# Whole Brain Cellular Activation After Mu and NOP Receptor Agonism: Shared Regional Targets with Divergent Network Consequences

**DOI:** 10.64898/2026.09.22.753526

**Authors:** Madeline Martinez, Akihiko Ozawa, Daniel Van Zant, Jake Thornberry, Lawrence Toll

**Author notes:** Corresponding author: Lawrence Toll, Stiles-Nicholson Brain Institute, Florida Atlantic University, 5353 Parkside Drive, Jupiter, FL 33458, USA.

## Abstract

**Rationale:** Mu and nociceptin/orphanin FQ peptide (NOP) receptors are both Gi/o-coupled and inhibitory, yet opioid agonists increase neuronal activation across the brain. How this occurs, and whether it differs between naive and dependent animals or between receptor subtypes, is unresolved.

**Objectives:** To map and directly compare whole-brain cellular activation after acute mu agonist treatment, in opioid-dependent animals and after NOP receptor agonism.

**Methods:** TRAP2/Ai9 reporter mice received acute morphine (10 mg/kg), an escalating twice-daily morphine regimen producing dependence, or the NOP agonist Ro 64-6198 (0.6 mg/kg). Brains were cleared and imaged intact by light-sheet microscopy; tdTomato-positive cells were quantified across 198 grey-matter regions registered to the Allen CCFv3, and interregional co-activation networks constructed.

**Results:** All treatments shifted activation upward relative to vehicle, significantly so for morphine in males; acute morphine engaged the most regions, all increased. Sex differences were confined to magnitude: the whole-brain increase was significant in males but not females, and no region showed a condition × sex interaction after correction. Ro 64-6198 produced a substantially weaker profile, smaller in magnitude rather than anatomically restricted, with no region surviving correction. Fewer regions were activated in dependent animals than after acute morphine, with no region distinguishing the two. Dependence substantially altered interregional co-activation and changed hub composition, while global network organization showed no detected change.

**Conclusions:** Mu and NOP agonism are distinguished by response magnitude and network role rather than by regional identity; dependence likewise acts principally on the coordination of activity rather than on which regions are recruited.

## Introduction

Opioid receptors are found throughout the brain and spinal cord, as well as in the periphery, consistent with the wide range of biological and physiological processes opioids influence, including pain, motivation, stress, respiration, and gastrointestinal motility (Mansour et al. 1995; Erbs et al. 2015; Ozawa et al. 2015). The three classical opioid receptors—mu, kappa, and delta—together with the nociceptin/orphanin FQ peptide (NOP) receptor make up the opioid receptor family, a group of Gi/o-coupled receptors activated by known endogenous peptides (Waldhoer et al. 2004; Le Merrer et al. 2009; Cox et al. 2015); yet nearly all clinically used medications are mu agonists. Morphine and analogs, through activation of mu receptors, are the most successful compounds for moderate-to-severe pain, but activation of these same receptors leads to serious complications including constipation, respiratory depression, reward and addiction (Koob and Volkow 2016). Agonists and antagonists of other receptors in the family have been developed for treatment of pain and other disorders but have been generally unsuccessful. Recently, the NOP receptor has drawn interest as a target for analgesia and substance-use medications, because NOP activation modulates reward processing and can temper the abuse liability and side-effects of mu agonists (Toll et al. 2016, 2019). As bifunctional NOP/mu agonists such as cebranopadol advance toward clinical use, a clearer understanding of how NOP receptor activation alters activity across the brain becomes increasingly valuable (Raffa et al. 2017; Coluzzi et al. 2022). We therefore included a selective NOP agonist alongside morphine to ask whether a second opioid-family receptor, signaling through the same Gi/o mechanism but with a different anatomical distribution, produces a comparable brain-wide activation signature to mu agonism, or a distinguishable one. Morphine was given both acutely and on an escalating regimen because tolerance and dependence are central to the clinical liability of mu agonists; NOP agonists are of interest precisely because they lack comparable dependence liability, so an equivalent escalating NOP regimen would not model the same process and was not included.

One limitation in understanding effects of acute and chronic opioid treatment has been inability to quantify cellular activation across the whole brain. This matters given the wide distribution of opioid receptors, whose activation alters activity at cellular, regional, and network levels. Opioid effects at the network level have been examined using fMRI in both humans and rodents, providing considerable information about resting state and drug-evoked functional connectivity among brain regions (Becerra et al. 2006; Sourty et al. 2024); however, fMRI lacks the resolution to investigate changes in activation at the cellular level. At cellular resolution, the primary approach has been to visualize the immediate early gene product Fos as a proxy for recent neuronal activity, including at the whole-brain level (Cai et al. 2022). More recently, mouse genetic models, such as TRAP (Targeted Recombination of Active Populations), in which CreER recombinase is expressed at the c-fos locus, and TRAP2/Ai9 mice, in which tamoxifen drives permanent tdTomato expression in neurons active during a defined window, have been used extensively (Madisen et al. 2010; Corder et al. 2019; DeNardo et al. 2019; Targowska-Duda et al. 2024; Wojick et al. 2025). This approach allows for a permanent label during a specific time window, but also further genetic manipulation due to the presence of Cre deposited in the activated neurons. The same strategy has been used to identify neurons activated by a discrete stimulus and then manipulate them with DREADDs (Targowska-Duda et al. 2024) or optogenetics (Allen et al. 2017).

Recent advances in optical clearing, light sheet microscopy, and automated cell counting have made it feasible to quantify tdTomato-labeled neurons throughout the brain. Because TRAP2/Ai9 yields a bright, endogenous tdTomato signal, cleared tissue can be imaged directly, without immunolabeling on which whole-brain Fos mapping has relied (Kimbrough et al. 2020; Ardinger et al. 2024; Vasylieva et al. 2026), thus removing any dependence on antibody selectivity or penetration into deep and internal structures. In our hands, whole-brain c-Fos immunolabeling gave inconsistent staining between brains together with an intense rim of signal at the brain surface arising from antibody accumulation. The resulting uneven signal intensity across the volume is poorly suited to automated, atlas-registered cell detection, which assumes comparable detectability throughout the brain. In this manuscript we use light sheet microscopy following clearing with a modified version of iDISCO adapted to preserve native fluorescence (MDISCO; Martinez et al. 2026b) to quantify neuronal activation throughout the mouse brain after treatment with acute morphine, escalating morphine doses to induce dependence, the selective NOP agonist Ro 64-6198, and vehicle. When registered to the Allen Common Coordinate Framework (CCFv3; Wang et al. 2020), sex-dependent changes in individual brain regions and functional co-activation were identified, with altered numbers of activated cells after drug treatment, while global network topology and modular organization were largely preserved, consistent with known behavioral, physiological, and neuroimaging observations in humans.

## Materials and Methods

All animal procedures were approved by the Florida Atlantic University Institutional Animal Care and Use Committee (IACUC; protocol no. A24-06) and conducted in accordance with the NIH Guide for the Care and Use of Laboratory Animals; the study is reported in accordance with the ARRIVE guidelines. Before treatment, all mice were habituated for 4 days to the testing environment, acrylic cylinders on a von Frey table, with daily saline i.p. injections; on the treatment day they remained in the cylinders from drug injection until return to the home cage, approximately 3.5 h, to limit non-drug mediated c-Fos activation. Male and female TRAP2 (Fos2A-iCreER) × Ai9 (Rosa26-LSL-tdTomato) mice (n = 6 per sex per condition, except Morphine-Dependent, n = 5) were randomly assigned to four conditions: Vehicle, Acute Morphine (10 mg/kg, i.p.), Morphine-Dependent (an escalating twice-daily morphine regimen, 10 to 75 mg/kg across Days 1–4 followed by a single 25 mg/kg injection on Day 5), or the nociceptin/orphanin FQ (NOP) receptor agonist Ro 64-6198 (0.6 mg/kg, i.p.; Mudgal et al. 2025). This dose of Ro 64-6198 is antinociceptive in a migraine pain model without producing sedation or motor impairment (Mudgal et al. 2025); higher doses are sedative (Morairty et al. 2023), which would itself alter brain-wide activity and confound the comparison with morphine. The escalating-dose regimen of morphine administration represents an established method to induce opioid physical dependence in rodents (Wilson et al. 2021); the reduced group size for this condition (n = 5 per sex) reflects loss of animals during the high-dose phase. Group sizes were chosen to match published whole-brain c-Fos mapping studies of similar design (Kimbrough et al. 2020; Ardinger et al. 2024; Vasylieva et al. 2026) rather than by a formal power analysis. Neurons active during drug exposure were labeled genetically rather than by immunostaining: 4-hydroxytamoxifen (50 mg/kg, i.p.), administered 30 min after the drug, drove permanent c-Fos–dependent tdTomato expression, and mice were perfused 10 days later. Whole brains were cleared with MDISCO (Martinez et al. 2026b), imaged intact on a Miltenyi UltraMicroscope Blaze II light-sheet microscope and processed in NeuroInfo (Tappan et al. 2019) for automated detection of tdTomato somata and registration to the Allen CCFv3 (Wang et al. 2020; Supplementary Methods; Online Resource 1). Image processing in NeuroInfo, including cell detection, atlas registration and region clean-up, was performed blind to condition; condition labels were unmasked only for statistical analysis. Automated detection and atlas-registration pipeline have been validated against manual delineation and reference-atlas landmarks (Eastwood et al. 2019; Tappan et al. 2019), with localization accuracy lower for small ventrolateral and midline structures. The complete experimental and analytical workflow is summarized in **Figure 1a–c**. Counts were restricted to grey-matter regions and averaged across hemispheres; bilateral sums were used for the negative binomial model, which requires integer counts. Regions were then filtered to remove the ventral-most structures obstructed at the imaging mounting surface and required to have bilateral representation in every export (16 further regions were absent or unilateral and were therefore not analyzed), yielding 198 regions for analysis (Table S1; Online Resource 2). Regions were analyzed at a single, consistent level of the CCFv3 summary-structure hierarchy, so that no analyzed region is nested within another and interregional co-activation is not inflated by anatomical containment.

**Fig. 1.**
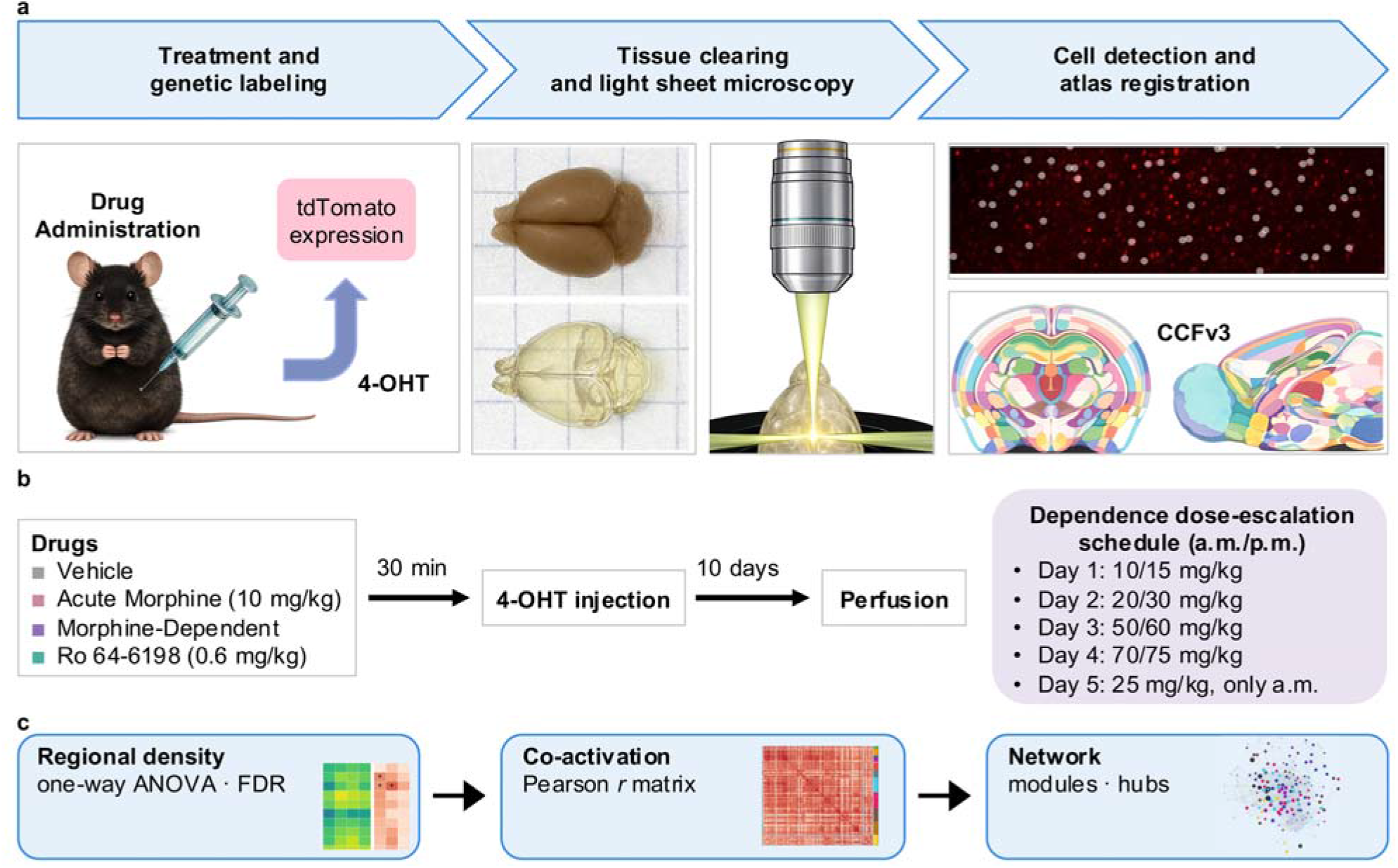
Experimental design and analysis workflow. (**a**) Whole-brain cellular activity mapping pipeline. TRAP2/Ai9 mice (Fos2A-iCreER × Rosa26-LSL-tdTomato) received drug treatment followed by 4-hydroxytamoxifen (4-OHT; 50 mg/kg, i.p.), permanently labeling neurons active during the labeling window with tdTomato. Brains were cleared with MDISCO, a modified iDISCO protocol adapted to preserve native fluorescence, and imaged intact by light-sheet fluorescence microscopy (Miltenyi UltraMicroscope Blaze II). tdTomato-positive cells were detected and registered to the Allen Mouse Brain Common Coordinate Framework (CCFv3), yielding cell counts and densities across 198 grey-matter regions. Left, drug administration and 4-OHT-dependent tdTomato labeling. Center, uncleared (top) and cleared (bottom) brain. Right, illustrative example of tdTomato signal in a morphine-dependent male with NeuroInfo-detected somata overlaid (grey), and the CCFv3 coronal and sagittal reference atlas. (**b**) Treatment timeline. Mice received vehicle, acute morphine (10 mg/kg, i.p.), morphine on an escalating dependence-inducing regimen, or the selective nociceptin/orphanin FQ peptide (NOP) receptor agonist Ro 64-6198 (0.6 mg/kg, i.p.), followed 30 min later by 4-OHT to open the labeling window. Brains were collected 10 d later. Right, twice-daily (a.m./p.m.) dose-escalation schedule for the morphine-dependent group, culminating in a single 25 mg/kg challenge on the morning of Day 5. Group sizes: Veh, Mor, and Ro, n = 12 (6 per sex); MorDep, n = 10 (5 per sex). (**c**) Analysis pipeline. Regional tdTomato-positive cell density was compared across conditions by one-way ANOVA on log10-transformed density, with Benjamini-Hochberg FDR correction applied across all 198 regions. Interregional co-activation was quantified as the Pearson correlation (r) between log10 regional tdTomato-positive cell counts across animals within each condition. Correlation matrices were thresholded at 10% matched density and analyzed as graphs to characterize modular organization and hub structure

Regional activity was analyzed as c-Fos density (cells/mm³, log_10_-transformed). The primary test was a one-way ANOVA across conditions (sexes pooled) with Tukey contrasts and Benjamini-Hochberg FDR across regions (q < 0.05); sex-stratified and condition × sex two-way ANOVAs were secondary and supplemental analyses, and a negative binomial GLM on bilateral summed counts, with log(region volume) as an offset, confirmed that primary findings were not an artifact of the log_10_ transformation. Interregional co-activation networks were constructed from across-animal Pearson correlations of log10 regional cell counts (combined, sex-residualized network as primary; per-sex networks reported as supplemental confirmation) and matched to a common 10% edge density. Networks were modular relative to degree-preserving nulls and small-world; condition differences in co-activation were tested edge-wise by Fisher r-to-z with FDR correction, and differences in global network metrics by bootstrap confidence intervals and label-permutation tests (strength and topology corrected as separate families). As a sensitivity analysis, regional densities were also re-analyzed after expressing each region relative to that animal’s whole-brain activation (global normalization; Supplementary Methods and Supplementary Results). Detailed procedures for clearing, imaging, registration, normalization, network construction, node-role assignment, and statistics are provided in Supplementary Methods (Online Resource 1). Effect sizes quoted for individual regions are those of regions selected as most extreme among 198 and are therefore subject to selection bias; they describe the ranked regions rather than providing unbiased estimates of the underlying effect.

Generative AI was used during manuscript preparation and in preparing the analysis code for public deposit. A detailed description of AI use can be found in Supplementary Information.

## Resources

Animals: male and female TRAP2/Ai9 mice (*Mus musculus*), bred in-house by crossing TRAP2 (Fos2A-iCreER; The Jackson Laboratory, stock #030323) with Ai9 (Rosa26-LSL-tdTomato; The Jackson Laboratory, stock #007909) mice, aged 6–8 weeks at the start of the experiments. Test compounds: morphine sulfate and Ro 64-6198 (98% purity) were supplied by the NIDA Drug Supply Program and dissolved in vehicle (5% DMSO, 0.5% hydroxypropyl cellulose); the Ro 64-6198 dose was chosen from published dose–response data as antinociceptive without sedation (Morairty et al. 2023; Mudgal et al. 2025); 4-OHT (Thermo Fisher Scientific) was given in corn oil. Instruments and software: UltraMicroscope Blaze II light-sheet microscope and MACS iQ View (Miltenyi Biotec); Imaris (version 11.1.0; Oxford Instruments); NeuroInfo (version 2026.1.1; MBF Bioscience); the Allen Mouse Brain Common Coordinate Framework v3 (Wang et al. 2020); R 4.4.3, with the packages listed in Supplementary Methods (Online Resource 1); analysis code version 1.1.0 (Martinez et al. 2026a). No antibodies or cell lines were used.

## Results

### Acute morphine engages a hypothalamic–preoptic–brainstem network

Regional activation across all 198 analyzed regions and all four conditions (vehicle, acute morphine, morphine-dependent, and Ro 64-6198) is shown in **Figure 2**, with the strongest pooled effects in **Figure 3**. When sexes were combined, acute morphine produced a significant 1.83-fold increase in global activation — quantified as each animal’s mean log10 regional c-Fos density across the 198 analyzed regions — relative to vehicle (one-way ANOVA F(3,42) = 5.97, p = 0.002; Tukey Vehicle vs. Morphine p = 0.001; Figure 4b). c-Fos density was also elevated in dependent animals (1.58-fold; Tukey p = 0.026), whereas this was not the case after Ro 64-6198 treatment (1.41-fold; Tukey p = 0.11). The effect on dependent animals was less robust than after acute morphine treatment: significant on mean regional density but weaker on whole-brain total cell counts (Tukey p = 0.056 for acute morphine and p = 0.31 for dependence; Supplementary Methods).

**Fig. 2.**
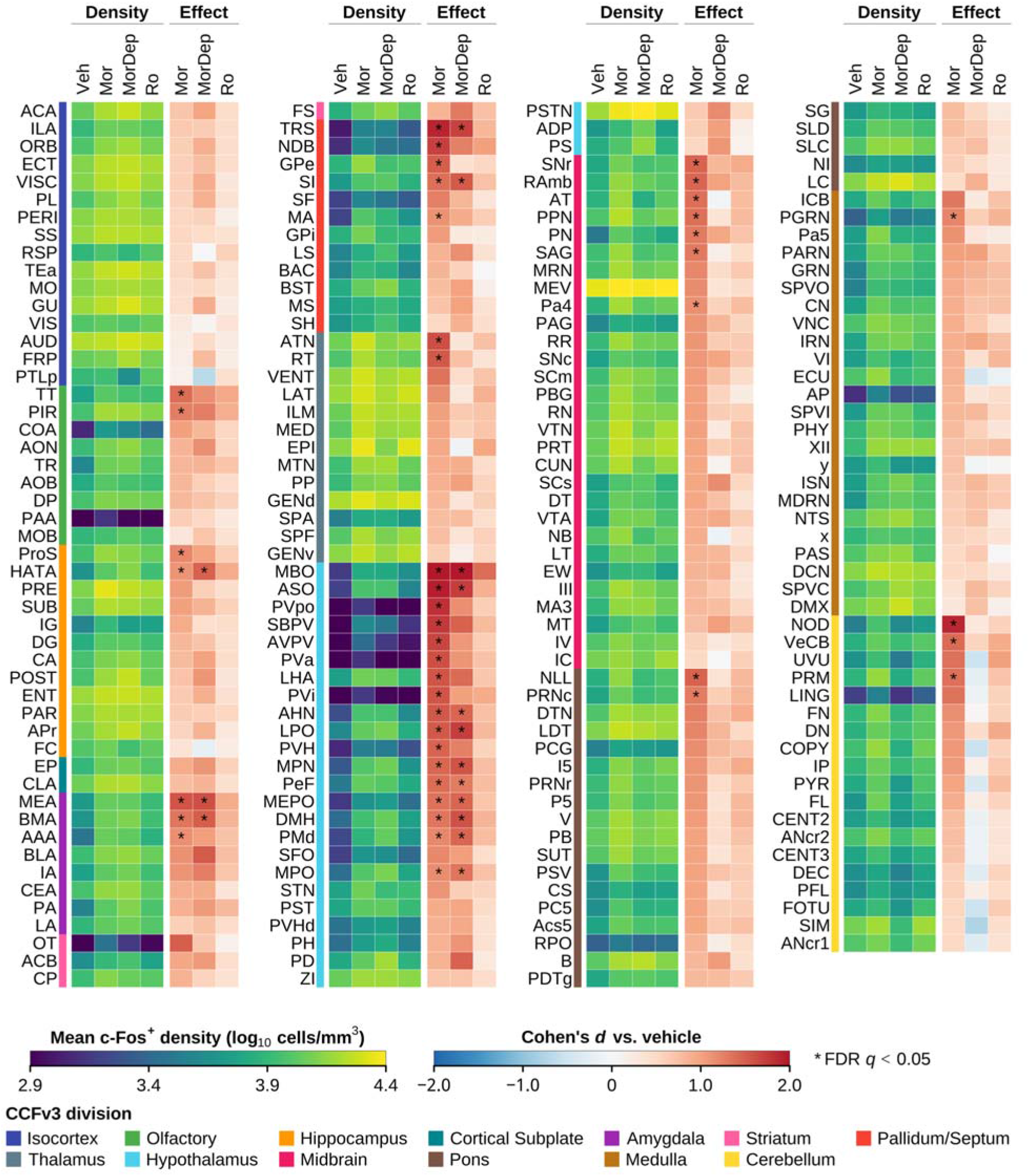
Brain-wide regional activation across all four conditions. Mean c-Fos density and effect size for each of the 198 analyzed grey-matter regions. Regions are split into four blocks of up to 50 rows to fit the page; blocks read left to right and the ordering runs continuously across them. Within a block each row is one region, labeled by acronym and tagged with a color band giving its CCFv3 division (key below). Rows are ordered by division, and by acute-morphine effect size within division. The left sub-block (Density) gives mean log10 c-Fos density per condition (Veh, vehicle; Mor, acute morphine; MorDep, morphine-dependent; Ro, Ro 64-6198) on a sequential scale; the right sub-block (Effect) gives Cohen’s d versus vehicle for each drug condition on a diverging scale centered on zero. Asterisks mark contrasts surviving Benjamini–Hochberg FDR correction across all 198 regions (q < 0.05): 44 regions under acute morphine, 15 under dependence, and none under Ro 64-6198. Both color scales are clipped at the 2nd and 98th percentiles of their distributions (density 2.9–4.4 log10 cells/mm³; effect ±2.0), because a few extreme regions would otherwise compress the figure into a narrow band of color; values beyond the limits saturate rather than being hidden, which affects 18 of 1386 cells (1.3%). Region acronyms and full names are given in Table S1, and per-region statistics in Table S2. Combined-sex n = 12/12/10/12

**Fig. 3.**
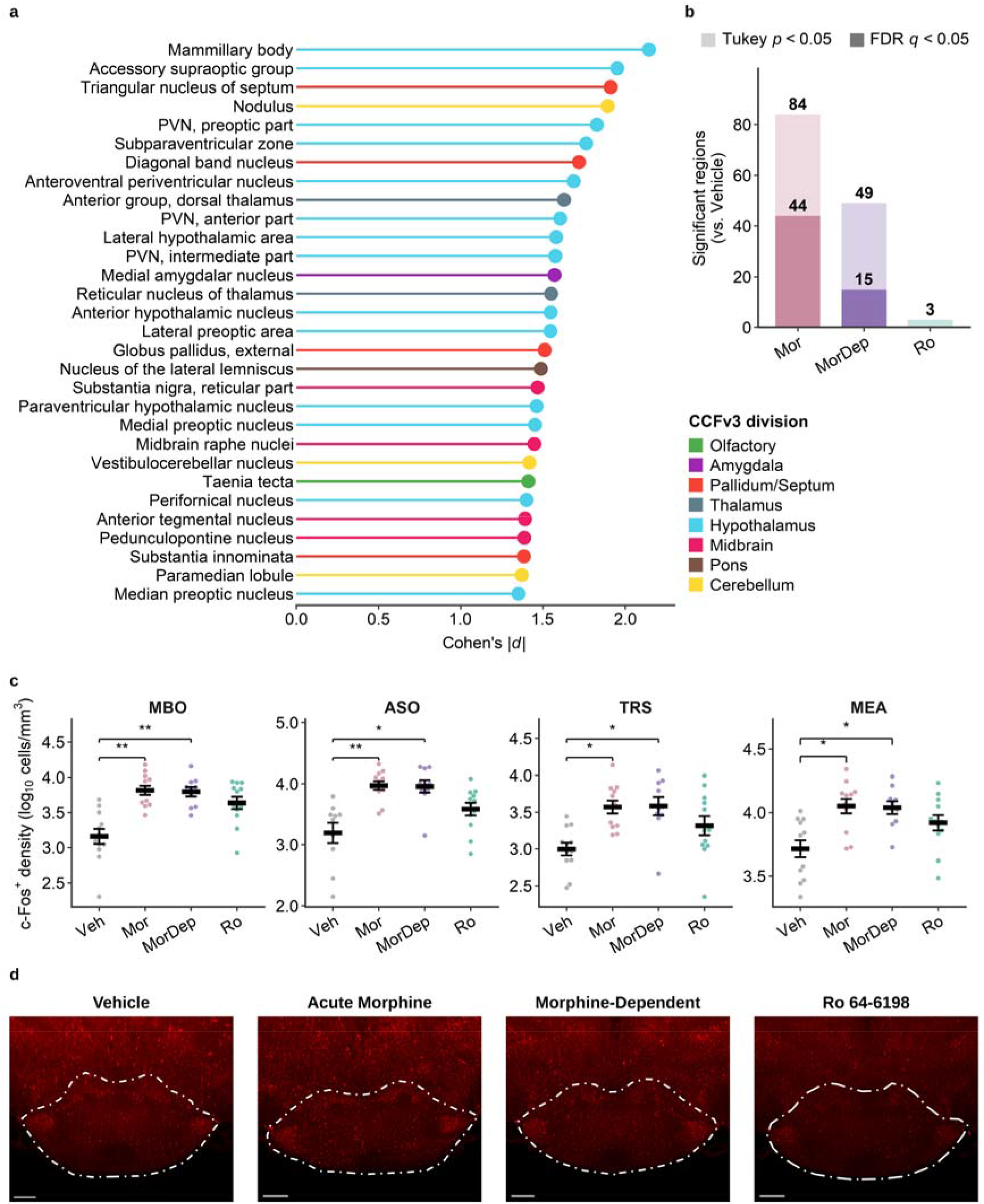
Whole-brain regional activation after opioid and NOP receptor agonists (combined sexes). (a) Lollipop plot of the 30 top-ranked regions by acute-morphine effect size (Cohen’s |d| vs. vehicle), ordered and colored by CCFv3 division. (b) Number of regions differing from vehicle for each treatment at Tukey p < 0.05 (light) and after Benjamini–Hochberg FDR correction, q < 0.05 (dark). (c) Exemplar c-Fos density (log10 cells/mm³) for the mammillary body (MBO), accessory supraoptic group (ASO), triangular nucleus of septum (TRS), and medial amygdala (MEA); points are individual animals, mean ± SEM; brackets show Tukey-adjusted comparisons vs. vehicle (*q < 0.05, **q < 0.01). (d) Representative light-sheet images of tdTomato (c-Fos) signal in the mammillary body (MBO, outlined; quantified in c) from one male per condition. Images are 40 µm optical sections rendered in Imaris with acquisition and display settings standardized across conditions within the application rather than under a single fixed display range, so they illustrate the labeling qualitatively and are not a quantitative intensity comparison; because a 40 µm section samples a fraction of the tissue volume over which NeuroInfo counts somata, the four conditions appear more similar than the counted difference. MBO c-Fos density under acute morphine was 3.88-fold that of vehicle in males (Cohen’s d = 2.29, FDR q = 0.040) and 3.88-fold in females (d = 1.92, q = 0.70). Scale bar, 500 µm. Combined-sex n = 12/12/10/12 (Vehicle/Acute Morphine/Morphine-Dependent/Ro 64-6198)

**Fig. 4.**
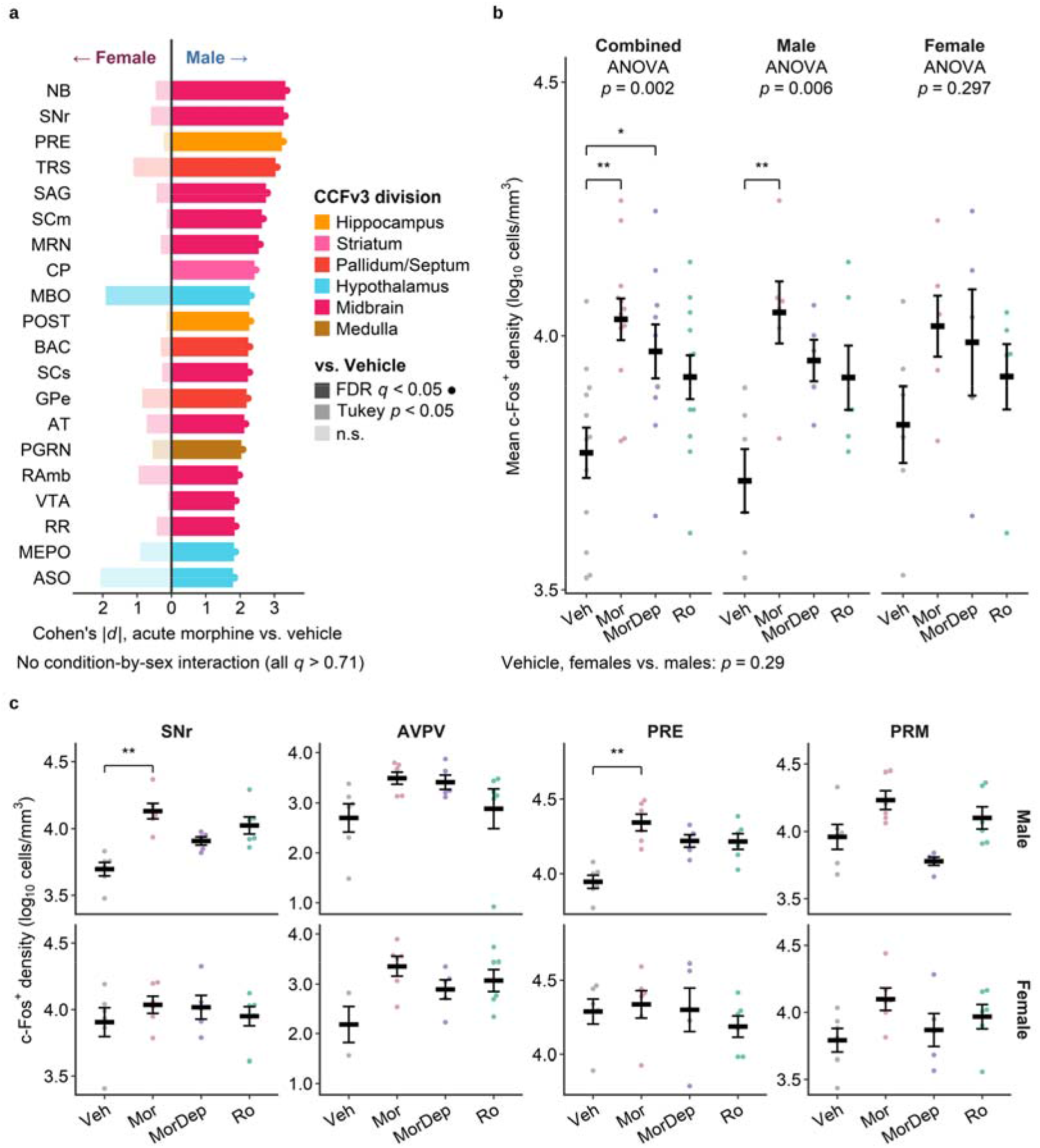
Regional activation differs in magnitude by sex and is largely sustained under dependence. (**a**) Diverging bar plot of sex-stratified effect sizes (Cohen’s |d| vs. vehicle) for the 20 regions reaching FDR q < 0.05 in either sex for the acute morphine versus vehicle contrast (the same criterion applied to both sexes), males (right) vs. females (left), ordered by male |d| and colored by CCFv3 division. Fill opacity indicates the significance tier within that sex — solid, FDR q < 0.05; mid, Tukey p < 0.05 uncorrected; faint, not significant — and a filled dot at the bar tip marks the FDR-significant bars. Region acronyms are defined in Table S1. No condition-by-sex interaction survived correction in any region (two-way ANOVA, all q > 0.71 across the 198 analyzed regions; per-region terms in Table S2). (**b**) Global activation (per-animal mean regional c-Fos density, log10 cells/mm³) per condition, shown pooled across sexes (Combined) and separately for males and females; each point is one animal, bars are mean ± SEM (one-way ANOVA: combined p = 0.002, males p = 0.006, females p = 0.297). Brackets show Tukey post hoc contrasts vs. vehicle and are drawn only within a stratum whose omnibus ANOVA is significant (*p < 0.05, \*\***p < 0.01,** ***p < 0.001). Vehicle males vs. females, p = 0.29. (**c**) Exemplar c-Fos density (log10 cells/mm³) for substantia nigra reticular part (SNr), anteroventral periventricular nucleus (AVPV), presubiculum (PRE), and paramedian lobule (PRM), split by sex across conditions; points are animals, mean ± SEM; brackets mark contrasts surviving FDR correction within that sex (*q < 0.05, **q < 0.01). AVPV and PRM are included as the exemplars discussed in the text — the largest female effect and the male cerebellar decrease respectively — and neither reaches FDR significance in either sex, so both are shown unbracketed. Per-sex n = 6/6/5/6

Across the four conditions, 84 regions showed elevated c-Fos density under acute morphine relative to vehicle at Tukey p < 0.05 (all Vehicle < Morphine), of which 44 survived Benjamini– Hochberg correction (q < 0.05; Table S2). The strongest effects were concentrated in the periventricular hypothalamus, preoptic area, and adjacent basal forebrain (all |d| > 1.5, **Figure 3a**), led by the mammillary body (MBO; F(3,42) = 13.11, p = 3.5 × 10; Tukey Vehicle vs. Morphine q = 0.002, |d| = 2.14, exemplified in **Figure 3c**) and including the accessory supraoptic group, periventricular and subparaventricular nuclei, diagonal band nucleus, and medial and lateral preoptic areas (**Figure 3a**). A secondary cluster spanned midbrain raphe, lateral lemniscus, and substantia nigra reticulata, with limbic extension through medial and basomedial amygdala and piriform cortex. The full ranked set, with FDR q-values, is reported in Table S2. Refitting the combined-sex comparisons with a negative binomial GLM on bilateral summed counts replicated 43 of the 44 FDR-significant regions under acute morphine, with 99.5% sign agreement across all 198 regions (e.g., MBO q = 2.2 × 10), confirming the effects were not an artifact of the log transformation. Despite morphine’s Gi/o coupling, none of the 198 regions showed a significant decrease in tdTomato expression. A breakdown of the number of significant and FDR-significant differences from vehicle for each treatment group is shown in **Figure 3b**. Following four days of escalating morphine (**Figure 1b**), the combined-sex analysis identified 49 regions at Tukey p < 0.05 versus vehicle, of which 15 survived FDR correction (Table S2). The periventricular-preoptic core identified after acute morphine treatment persisted in dependent animals — MBO (Tukey Vehicle vs. Morphine-Dependent q = 0.008, |d| = 2.06), accessory supraoptic group, medial preoptic and anterior hypothalamic nuclei (all FDR-significant). Every region significant under dependence was also significant under acute morphine: rather than recruiting a distinct regional profile, the 15 FDR-significant regions formed a strict subset of the 44 engaged acutely, and no region crossed the FDR threshold uniquely under dependence. Additional forebrain and striato-limbic regions (lateral septum, accumbens, basolateral amygdala, anterior cingulate) reached significance (Tukey p < 0.05) only, accompanying the persistent FDR-significant core (Table S2). These pooled results nonetheless obscured a substantial difference in the magnitude of the response between the sexes.

### Morphine-induced activation is larger and more consistent in males

Morphine-induced activation was in the same direction in both sexes but differed in breadth and in the magnitude reached, the increase being larger and more consistent in males. This asymmetry is apparent in global activation: mean regional c-Fos density rose 2.15-fold in males (one-way ANOVA F(3,19) = 5.77, p = 0.006; Tukey Vehicle vs. Morphine p = 0.004) and 1.56-fold in females, where it did not reach significance at this sample size (F(3,19) = 1.32, p = 0.30) (**Figure 4b).** Both sexes therefore moved in the same direction and the pooled increase (Figure 4b) is the sum of a significant male and a non-significant female response; the condition × sex interaction in global activation was not significant (F(3,38) = 0.40, p = 0.75), so the sexes are not formally distinguishable on this measure. In within-sex ANOVAs, males engaged a broad set of 82 regions (20 FDR-significant despite n=6 per group rather than n=12 in the pooled analysis), with the most highly activated regions encompassing midbrain structures, together with collicular, septal, basal ganglia, VTA, descending pain circuitry (PAG, PPN, RN) and a distinctive hippocampal-subicular cluster (presubiculum, postsubiculum, dentate gyrus, CA fields), none of which were detected in females (Figure 4a). This wider male set may reflect the larger male global response rather than a distinct male circuit: none of the 20 FDR-significant male regions retains FDR significance when each region is expressed relative to that animal’s brain-wide activation. Under that alternate normalization the pattern reverses, with the number of regions at uncorrected p < 0.05 falling from 82 to 13 in males and rising from 9 to 24 in females (Supplementary Results). Top male effects exceeded |d| = 2.5, exemplified by the substantia nigra reticular part and presubiculum (SNr and PRE; **Figure 4c; Table S2; Online Resource 3)**. Females reached significance (Tukey-adjusted p < 0.05, not corrected across regions) in only 9 regions, none surviving FDR, concentrated in preoptic and periventricular hypothalamic targets. Despite the lack of FDR significance, the effect level in several hypothalamic regions was quite large, with the anteroventral periventricular nucleus showing the largest female effect (AVPV; |d| = 2.22, well above its combined-sex estimate, **Figure 4c**). Because regions are correlated, counts at this threshold were calibrated by permuting condition labels within sex (10,000 permutations; null median 0 regions, 95th percentile 11 in each sex). The male count far exceeds this null (p < 0.001), whereas the female count of 9 regions does not (p = 0.062), so the large individual female effects, such as the AVPV, are reported region by region rather than as evidence of a female-specific regional pattern. Two-way ANOVA revealed no condition × sex interaction surviving FDR at any region (q > 0.71; 12 nominal at uncorrected p < 0.05). Baseline activation also differed numerically between sexes: vehicle-treated females showed 1.29-fold higher activation than vehicle-treated males, higher in 144 of 198 regions, although this difference was not significant at the present sample size (Welch t-test on vehicle animals, p = 0.29). A higher baseline compresses the achievable fold-change and is consistent with the smaller female response being one of magnitude rather than of regional pattern.

### Morphine dependence sustains the hypothalamic–preoptic activation core

Dependence sustained the pooled activation core rather than recruiting new regions: 49 regions differed from vehicle at Tukey p < 0.05 and 15 survived FDR correction, and all 15 lie within the 44 regions identified under acute morphine, with no dependence-specific region anywhere in the brain (**Figure 2**). Across those regions the mean effect size fell from d = 1.47 under acute morphine to d = 1.05 under dependence, and the two regional profiles correlate at r = 0.35 across all 198 regions, so the core is retained at reduced amplitude. Dependent animals also showed sex differences in regional neuronal activation. Males showed 38 regions at Tukey p < 0.05 (none FDR-significant; p = 0.010 against the permutation null), with the largest effects extending beyond the core into septal, collicular, and midbrain nodes (triangular nucleus of septum |d| = 3.74) while the core itself remained strongly represented (Table S2). Females showed only 7 regions (none FDR-significant), again at preoptic-hypothalamic targets, although this count does not exceed the permutation null (p = 0.078) and is not interpreted as a distinct female footprint. Direct comparison of the two morphine conditions revealed a non-significant decrease in total activation and no region surviving FDR whether looking at pooled or individual sex data (0/198 FDR-significant), although a set of cerebellar, pontine, and medullary structures showed lower c-Fos density in dependent animals compared to acute morphine or vehicle treatment (Tukey p < 0.05), an effect driven entirely by males and exemplified by the paramedian lobule (**PRM; Figure 4c**). Dependence was also the only condition in which a substantial fraction of regions fell below vehicle (21 of 198, 10.6%; minimum d = −0.75), compared with none under acute morphine and one under Ro 64-6198. Accordingly, in dependent animals, regional effects were the most variable of the three conditions, with standard deviation of regional effect sizes 1.36 times that under acute morphine.

### NOP receptor agonism produces a weaker activation profile with a similar regional distribution

Acute Ro 64-6198 (0.6 mg/kg, analgesic but not sedative; Mudgal et al. 2025) reached significance (Tukey p < 0.05) in only 3 regions versus vehicle (MBO [**Figure 3d],** diagonal band nucleus, intertrigeminal nucleus), none surviving FDR — compared to 84 under acute morphine and 49 under dependence at the same threshold (**Figure 3b**; Table S2). This pooled count does not exceed the permutation null (p = 0.14), and these three regions are therefore not interpreted as specifically engaged. Effect sizes nonetheless remained positive in 99.5% of regions but smaller than morphine’s (mean |d| 0.56 vs 0.98), and the Ro 64-6198 regional effect profile correlated with the acute morphine profile at r = 0.63 across all 198 regions — more strongly than dependence did (r = 0.35) — with Ro effect sizes scaling at approximately 0.40 of the acute morphine profile. The NOP response is therefore weaker rather than anatomically restricted (**Figure 2**). Because all three profiles are referenced to the same vehicle group, these correlations are mutually comparable but are not independent estimates of profile similarity (Supplementary Methods; Supplementary Results). Sex-stratified analysis again showed the same asymmetry in detection: males engaged 18 regions (none FDR-significant; p = 0.027 against the null) overlapping the male brainstem-mesencephalic morphine signature (SAG, SNr, PRE, SCs, all |d| > 2.2), whereas females showed no region at Tukey p < 0.05. As with acute morphine, this male count falls sharply under global normalization (18 regions to 6; Supplementary Results), so it is read here as a difference in the magnitude of the response rather than in which regions are engaged. AVPV, the largest female node under acute morphine, showed a comparable female effect size under Ro 64-6198 (|d| = 1.56) but did not reach significance.

### Widespread increases in co-activation under morphine dependence occur without detected changes in global network organization

The regional and network analyses differed in what they detected: acute morphine produced the broadest change in individual-region activity, whereas dependence produced, by far, the largest change in inter-regional co-activation despite engaging fewer regions. To ask how drug treatment reorganizes coordination between regions, we built co-activation networks from across-animal correlations in regional activated-cell counts (log10 transformed; Pearson r; region pairs varying in tandem treated as ’co-activated’), pooling all animals after mean-centering each region within sex to remove overall male– female differences. Per-sex networks confirmed that combined-network results were maintained within each sex (Supplementary Methods). In the combined-sex data, mean absolute correlation across all 19,503 region pairs was lowest under vehicle (|r| = 0.45), intermediate after acute morphine (0.52), far higher in dependent animals (0.74), and near vehicle levels after Ro 64-6198 (0.50; **Figure 5a,b**). For topological comparisons, networks were thresholded to the strongest 10% of links (matched density; Rubinov and Sporns 2010). This rise in mean correlation was broad-based rather than system-specific, with within-division co-activation increasing in dependent animals in 11 of the 12 divisions shown, isocortex being the exception (**Figure 5d**). The increase in co-activation in dependent animals was present in both sexes, and numerically larger in females (**Figure 5b**; per-sex and sex-difference matrices, Figure S1a,c), although this excess does not survive removal of each animal’s global activation level (Figure S1d; Supplementary Results). Because correlation magnitude is sensitive to sample size and the dependent group had the smallest n, all conditions were subsampled to the dependent-group n (5 per sex, 2000 draws). Dependent-group co-activation (mean |r| = 0.74) exceeded the 97.5th percentile of every subsampled control distribution (vehicle 0.53, acute morphine 0.57, Ro 64-6198 0.59), excluding sample size as an explanation (Supplementary Results).

**Fig. 5.**
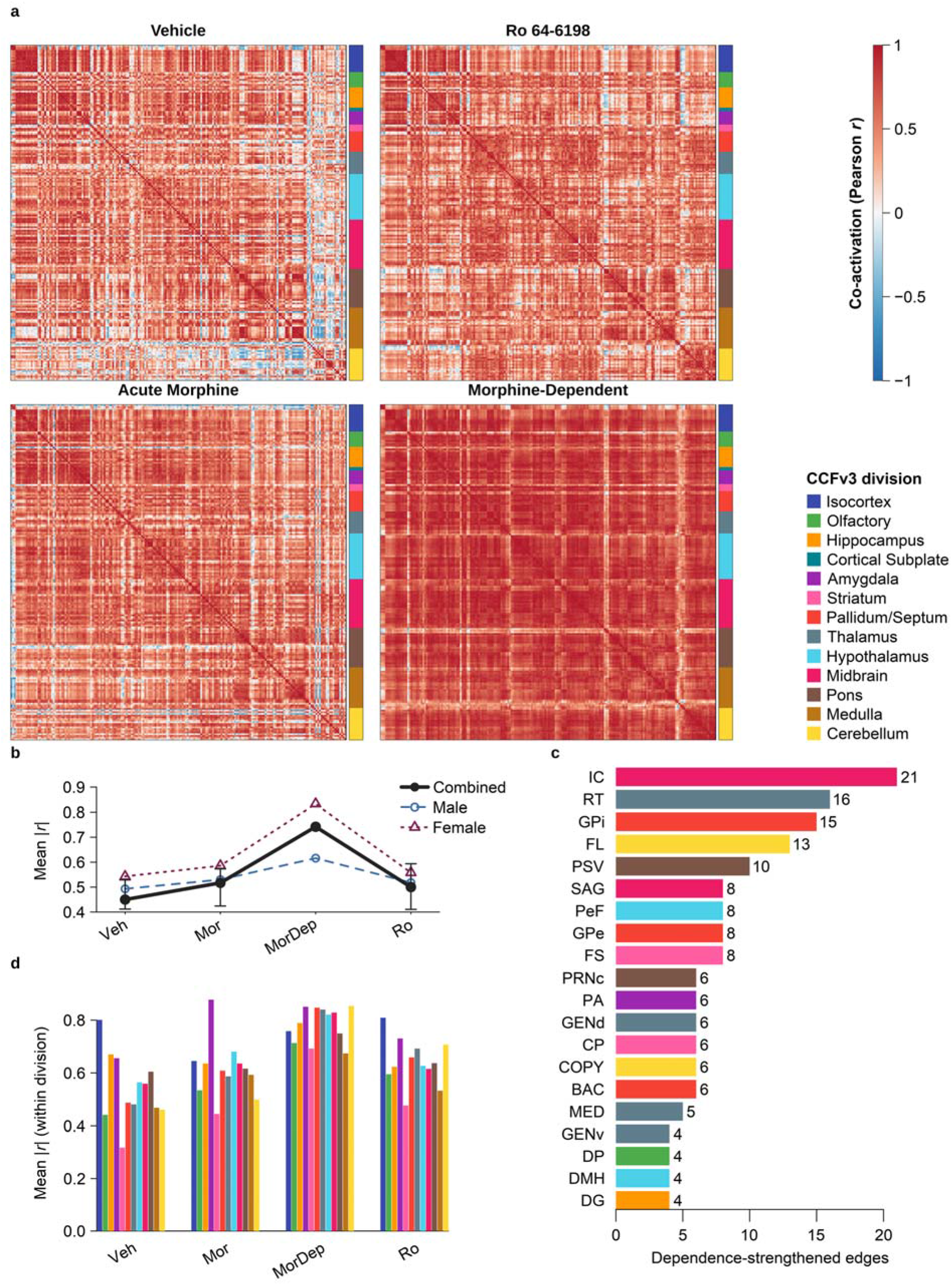
Altered brain-wide co-activation under morphine dependence. (**a**) Region × region co-activation matrices (Pearson r across animals, 198 grey-matter regions) for each condition (Vehicle and Ro 64-6198, top; Acute Morphine and Morphine-Dependent, bottom); regions ordered by CCFv3 division, indicated by the anatomy bar beside each matrix and named in the CCFv3 division key (right), with a shared correlation scale; all 198 regions are listed by abbreviation, full name and division in Table S1. (**b**) Mean absolute co-activation (mean |r|) per condition for the combined network and for each sex. Whiskers on the combined series show the 2.5th–97.5th percentile of mean |r| when every condition is subsampled to the smallest group n (5 per sex, 2,000 draws, the network rebuilt on each draw); the morphine-dependent group is already at that n and is plotted as a point. (**c**) Region involvement among the 160 region pairs with significantly strengthened co-activation under dependence (Vehicle vs. Morphine-Dependent; Fisher r-to-z, BH-corrected), shown for the 20 most-involved of the 101 regions represented and colored by division, with the number of pairs at each bar end. Because each pair involves two regions, bars sum to more than the number of pairs. The same comparison after removal of the per-animal global offset is shown in Figure S1d,e. (**d**) Mean within-division co-activation (mean |r|) by division and condition, bars colored by division as in the CCFv3 division key. The cortical subplate is omitted from **d** (2 regions form a single within-division pair, insufficient for a stable mean), but still appears in the heatmap anatomy bars and CCFv3 division key. Combined-sex n = 12/12/10/12

Edgewise comparison (Fisher r-to-z, BH-corrected) localized these differences in co-activation strength almost entirely to dependence, as the vehicle versus morphine-dependent comparison yielded, in the combined-sex network, 160 region pairs with significantly altered co-activation, every one of them stronger in dependent animals. Every other pairwise comparison among the four conditions (e.g., vehicle vs. acute morphine, vehicle vs. Ro 64-6198, acute vs. dependent) yielded ≤ 3 such pairs (**Figure 5c; condition-difference matrices, Figure S1b; Table S3; Online Resource 4)**. A sensitivity analysis that removed each animal’s global activation level before computing correlations (global signal regression) is reported in Supplementary Results (Figure S1d,e).

### Hub composition differs across conditions while global network organization shows no detected change

Although dependence sharply increased co-activation strength, we detected no accompanying change in the network’s overall architecture — how it organizes into communities or how efficiently it is wired. In all four conditions, the networks shared two features typical of brain networks. They were modular — regions clustered into communities far more than in degree-matched random networks (z = 55.8–79.3 combined; z = 50.8–82.6 per sex; all p < 0.001) — and small-world, combining dense local clustering with short paths between distant regions (σ > 1 in all networks; Table S4; Online Resource 5). Summary measures of architecture were compared across conditions: node strength, transitivity, global efficiency, and modularity Q showed no detected differences across treatments (permutation tests, all q ≥ 0.237; overlapping bootstrap 95% CIs). The precision of this null varies by metric: with the present sample the bootstrap intervals exclude differences larger than approximately 10–17% of the vehicle value for node strength, transitivity and global efficiency, but only differences larger than approximately 37% for modularity Q, so equivalence should not be inferred for modularity. Global efficiency was the exception: it was 19% higher under dependence than under vehicle, the largest of these differences and slightly above the 16% resolvable for that metric. The permutation test nonetheless did not reject the null (p = 0.11, q = 0.34), so this difference is reported as undetected rather than absent, and a real change in global efficiency of this magnitude cannot be excluded. Thresholded correlation networks are expected to show elevated clustering and small-worldness relative to degree-preserving random graphs by construction, so these comparisons are reported as confirmation that the networks are well formed rather than as biological findings. The null used for these comparisons does not preserve the transitivity that correlation induces between region pairs, which inflates apparent small-worldness specifically; the comparisons are therefore descriptive and no condition contrast rests on them (Supplementary Methods).

A community-detection algorithm (consensus Louvain) split each combined-sex network into 5–8 modules (5–10 across the per-sex networks) that were stable across treatments. The module assignment of each region under each condition is reported in Table S5 (Online Resource 6). We summarized each region’s position in the network with two standard cartographic measures (Guimerà and Nunes Amaral 2005): the participation coefficient (PC), which is high when a region’s connections are distributed across multiple modules (a between-module ’connector’), and the within-module degree z-score (WMDz), which is high when a region is strongly connected within its own module (a ’provincial hub’). Their anatomical distribution across conditions is mapped for the combined network (**Figure 6a,b**) and for each sex, where the male and female maps diverge substantially (Figure S2a,b). The per-sex maps diverged most in isocortex, where acute morphine produced numerous high-PC connector regions in females but few in males, though the reduced per-sex sample precludes formal comparison. The joint PC–WMDz distribution used to classify cartographic roles is shown in Figure S3. The vehicle network was dominated by locally connected regions with few bridges (**Figure 6c**), with bridging ’connector’ regions (high PC) becoming more common after acute morphine treatment (median PC 0.20 vs. 0.43; **Figure 6a**); Ro 64-6198 treatment induced regions that were highly connected but mainly within their own module (high WMDz; ’provincial hubs’). Hub composition also differed. By the multi-centrality convergence criterion (agreement across four centrality measures; Supplementary Methods), anterior cingulate and prelimbic cortex were hubs in all four conditions, while several cortical, olfactory, and hippocampal hubs of the drug-naive network dropped out in dependent animals. In **Figure 6c**, rings mark all hub-role regions (WMDz ≥ 1.5) and labeled nodes are the connector-hub subset. Connector hubs under acute morphine were hypothalamic (lateral hypothalamic area, perifornical nucleus) together with midbrain and pontomedullary nodes; under dependence they were hypothalamic (dorsomedial and anteroventral periventricular nuclei), hippocampal, striatal (nucleus accumbens), and medullary. Per-sex networks confirmed this pattern but with less stable hub assignments (no region was a hub in all four conditions in males, and only anterior cingulate in females); female multi-centrality hub counts rose under each drug condition (24 under vehicle vs. 31–35 under morphine and Ro 64-6198) relative to the more stable male count (32–37). Hub assignment was assessed for stability by leave-one-animal-out jackknife, in which each network was rebuilt with each animal left out in turn and the hubs recomputed (Supplementary Results). The full-sample hub set was recovered with mean Jaccard indices of 0.56–0.82 across conditions; prelimbic cortex was recovered in 92–100% of jackknife networks in every condition, whereas anterior cingulate was recovered in 92–100% of vehicle, acute morphine and Ro 64-6198 networks but in only 50% of dependent ones, so its hub status under dependence should be regarded as unstable. Of the 30 regions that lost hub status in dependent animals, 24 were hubs in at least 10 of the 12 vehicle jackknife networks and in no more than 3 of the 10 dependent ones. Because jackknife replicates are not independent, these recovery rates quantify the stability of hub assignment rather than constituting a formal test of difference between conditions, and the change in hub composition is accordingly described rather than tested.

**Fig. 6.**
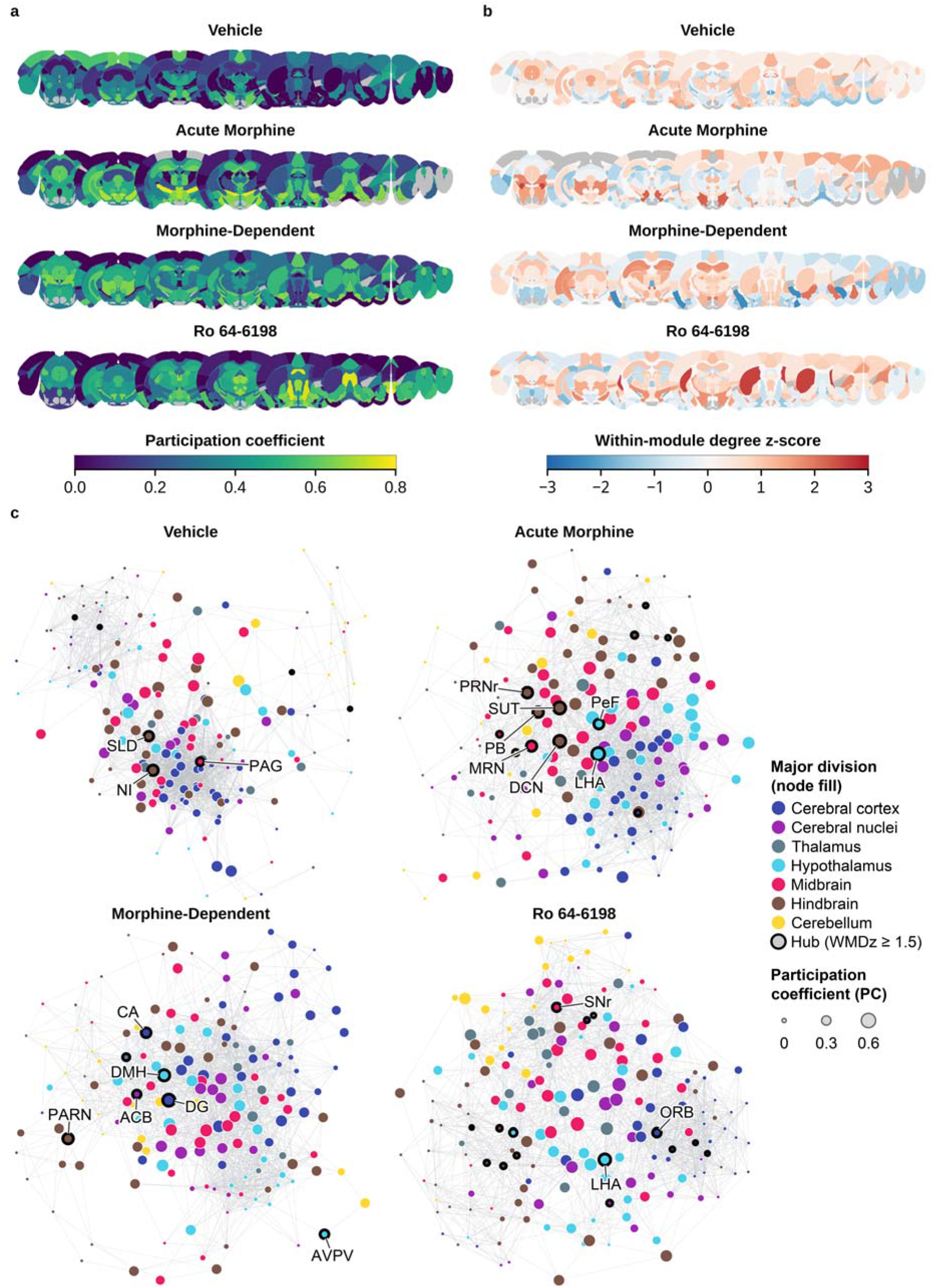
Network cartography and organization. (**a**) Participation coefficient (PC) mapped onto nine evenly spaced CCFv3 coronal levels for each condition (PC scale 0–0.8); regions with no value at that level are grey. (**b**) Within-module degree z-score (WMDz) on the same coronal levels and conditions (scale −3 to 3), grey as in **a**. (**c**) Force-directed (Kamada–Kawai) representations of each co-activation network (density-matched at 10% edge density, 1,951 edges per network); node fill indicates major brain division, node size scales with PC, and black rings mark hubs (WMDz ≥ 1.5); labeled nodes are connector hubs. Layouts are rescaled per network, so inter-node distances are not quantitatively comparable across conditions. For node fill, the 13 CCFv3 divisions are collapsed to 7 major divisions (Isocortex, Olfactory, Hippocampus, and Cortical Subplate → Cerebral cortex; Amygdala, Striatum, and Pallidum/Septum → Cerebral nuclei; Pons and Medulla → Hindbrain; Thalamus, Hypothalamus, Midbrain, and Cerebellum retained). Combined-sex networks, n = 12/12/10/12 (Vehicle/Acute Morphine/Morphine-Dependent/Ro 64-6198)

## Discussion

Like many GPCRs, mu opioid receptors are distributed throughout the brain, and like the other members of the opioid receptor family (kappa, delta, and NOP), they are Gi/o coupled, so that receptor activation lowers cAMP, increases GIRK (G protein coupled, inwardly rectifying potassium) channel conductance, and generally inactivates the cell (Williams et al. 2013). Despite this inhibitory coupling, morphine-mediated disinhibition of GABAergic neurons raises extracellular dopamine in the nucleus accumbens, producing reward and abuse liability (Wise and Bozarth 1985; Di Chiara and Imperato 1988; Spanagel et al. 1990; Johnson and North 1992), and when injected into the rostral ventromedial medulla (RVM) activates “off” cells in the descending pain modulation pathway producing analgesia (Moreau and Fields 1986; Fields 2004; Heinricher et al. 2009). Therefore, mapping where and how strongly neurons are activated is central to understanding opioid action. Such activation has most often been inferred from the immediate-early gene c-Fos, measured by in situ hybridization or immunohistochemistry (Morgan et al. 1987; Sagar et al. 1988; Bullitt 1990; Herrera and Robertson 1996). Although these techniques visualize neuronal activation, temporal uncertainties and antibody variability complicate interpretation. The recent utilization of TRAP2 and TRAP2/Ai9 mice addresses both limitations, providing a permanent, genetically encoded record of neurons active within a defined window after a known stimulus, together with genetic access to those neurons for subsequent manipulation (Madisen et al. 2010; DeNardo et al. 2019; Targowska-Duda et al. 2024).

Combined with advances in tissue clearing and light sheet imaging, the use of TRAP2/Ai9 mice has enabled comprehensive whole-brain analysis of state-dependent cellular and regional neuronal activation, neuronal ensembles and interregional co-activation (Vetere et al. 2017). fMRI has identified connectivity changes associated with opioid treatment and opioid use disorder in humans (Becerra et al. 2006; Sourty et al. 2024) but lacks the cellular resolution and experimental control achievable in mice.

Here, whole-brain activation was mapped in TRAP2/Ai9 mice across four conditions: vehicle, acute morphine, an escalated regimen producing dependence, and acute treatment with the NOP receptor agonist Ro 64-6198. One major finding was that both morphine and Ro 64-6198 shifted global activation upward, although the increase reached significance only for the morphine conditions and, within sex, only in males, with morphine being more effective than Ro 64-6198. This is consistent with NOP receptor agonist activity being more modulatory (Pan et al. 2000), with receptors on both GABAergic (to disinhibit) and dopamine neurons (to inhibit) in the VTA (Petrella et al. 2024) and on both analgesic “primary cells” and nociceptive “secondary cells” in the PAG (Vaughan et al. 1997), while mu receptors act on GABAergic interneurons within these circuits, contributing to disinhibition-mediated analgesia and reward (Johnson and North 1992; Vaughan et al. 1997). These studies are consistent with chronic treatment reducing the level of neuronal activation compared to a single morphine administration, in keeping with the microarray studies (Korostynski et al. 2007), likely due to tolerance development; in the present data, however, no specific region differed significantly between the two morphine conditions. Because testing for withdrawal would compromise the neuronal analysis, we did not independently confirm dependence prior to perfusion; however, multiple-day ascending doses have been routinely used to induce dependence. Because activation was captured within a single TRAP labeling window, these maps represent one temporal snapshot of drug action. This window, approximately 6 h initiated by the 4-OHT injection (DeNardo et al. 2019), is considerably broader than the 90-120 min post-stimulus interval typical of c-Fos immunostaining. Because mu and NOP receptors are both Gi/o-coupled and inhibitory at the cells expressing them, the brain-wide increases reported here are unlikely to reflect direct receptor-mediated excitation of the labeled neurons. They are more consistent with disinhibition and other circuit-level consequences, in which inhibition of GABAergic interneurons releases downstream populations from tonic suppression; thus activity-dependent labeling reports the global neuronal consequence rather than the site of receptor action. Two consequences follow from the labeling window itself. Activation is integrated rather than sampled, so sequential waves of activation cannot be resolved and regions active at different times within the window are labeled together. Transient activation early in the window is weighted equally with sustained activation throughout it.

A second major finding was a sex difference in the magnitude of the response to 10 mg/kg morphine. Acute morphine reached significance in many more brain regions in males than in females, but this difference in breadth may not be independent of the difference in global activation, which rose significantly in males but not in females, possibly in part because of the numerically lower baseline activation in males. Because regional density is analyzed in absolute terms, a larger whole-brain increase raises the effect size in every region simultaneously, and detection breadth follows. Consistent with this, when each region is expressed relative to that animal’s brain-wide activation as a sensitivity analysis, the male regional advantage does not persist (Supplementary Results). We therefore interpret the sex differences as reflecting mainly overall response magnitude, although a contribution of regional selectivity cannot be excluded. The pooled analysis after acute morphine treatment identified hypothalamic regions, but extensive midbrain, collicular, and hippocampal–subicular activation predominated in males, consistent with higher mu-opioid receptor density in the PAG of male rats (Loyd et al. 2008) and greater intracerebroventricular analgesic potency in male C57BL/6 mice (Kest et al. 1999). However, such sex differences in morphine analgesia are strain-and species-dependent, with no sex differences in receptor expression or systemic morphine-induced analgesia in this strain (Boorman et al. 2025). Female cellular activation was concentrated in preoptic and periventricular hypothalamic targets, with no increase in midbrain activity. This is consistent with morphine strongly engaging hypothalamic neuroendocrine circuits that are modulated by sex hormones, with estrogen modulating mu-opioid receptor signaling in these circuits (Lee and Ho 2013). Notably, no region showed a condition × sex interaction surviving correction, indicating that both sexes engaged the same network in the same direction, differing in the magnitude of recruitment rather than in opposing responses. A recent whole-brain map of acute morphine reached a convergent conclusion on direction, reporting larger male responses and more regions engaged in males (45 versus 10 at one hour post-injection), but did detect region-level condition × sex interactions (Vasylieva et al. 2026). The discrepancy is consistent with the labeling window: TRAP2 integrates activity over approximately six hours, whereas that study sampled c-Fos protein at fixed one-and four-hour points, and its own sex effects fell from 80 structures at one hour to 10 at four hours. A wider integrating window would be expected to attenuate localized sex differences further. A second whole-brain study using TRAP2-based tagging has since mapped morphine-administration and withdrawal ensembles and reported them to be largely non-overlapping at the cellular level (Ishii et al. 2025); the present dependence condition, labeled at a day-5 challenge rather than during precipitated withdrawal, is not directly comparable but converges with that work in showing that opioid state is represented brain-wide rather than within a restricted set of regions.

A third major finding emerged only when activity was examined across regions rather than within them. Whereas acute morphine produced the broadest change in the number of activated regions, it was morphine dependence that produced the largest change in the coordination of activity between regions. In dependent animals of both sexes, the average correlation in activation between region pairs rose well above vehicle, acute morphine, and Ro 64-6198 levels, and this difference was not attributable to the smaller dependent-group sample. Two components contribute: a uniform rise that tracks animal-level variation in overall activation, and a redistribution of which specific region pairs co-vary, which persists when the global component is removed. This separation suggests that dependence reorganizes the relationships among regions more than it changes which regions are recruited.

A rise in interregional co-activation with dependence is consistent with whole-brain studies of alcohol, in which dependence and abstinence likewise increase cross-correlation between regions (Kimbrough et al. 2020; Bloch et al. 2021; Ardinger et al. 2024). In those studies, the increase in coupling was accompanied by an apparent reorganization of network architecture, most often a collapse of modular structure. We did not observe such a change, and the difference is most likely methodological. In the previous studies, networks were built at a fixed correlation threshold, so a condition with higher overall correlation necessarily produces a denser graph, and density strongly shapes measures of modularity and efficiency. We held density constant across conditions, so that architecture could be compared independently of correlation strength. Under these conditions, the coarse organization of the network did not change. Across all four conditions the networks remained modular and small-world, and summary measures of how efficiently activity could spread, how strongly regions clustered, and how cleanly the network divided into communities showed no detected differences, within the precision set out in the Results.

What did change was the identity of the most highly connected hub regions. Anterior cingulate and prelimbic cortex served as hubs in every condition, but under morphine several cortical, olfactory, and hippocampal hubs of the drug-naive brain gave way to hypothalamic and amygdalar regions — the same territories that dominated the regional analysis. The network and regional results therefore converge on a common set of hypothalamic structures, with prominent midbrain engagement in males, as central to the morphine-activated brain. Amygdalar regions appear among the multi-centrality hubs (anterior amygdalar area under acute morphine, posterior amygdalar nucleus under dependence) but not among the cartographic connector hubs labeled in **Figure 6c**. Notably, these were not the regions most often associated with opioid reward or dependence: the nucleus accumbens and ventral tegmental area, and the central amygdala and bed nucleus of the stria terminalis, all shifted in the same direction as the surviving regions but did not themselves reach significance after correction in the pooled analysis. In males alone, the nucleus accumbens, VTA, and bed nucleus of the stria terminalis were nominally significant (Cohen’s d = 1.6–1.8) but were essentially absent in females (d < 0.4). What survived correction, given the limited sample, was a predominantly neuroendocrine and hypothalamic-integrative profile, comprising preoptic and hypothalamic nuclei together with the medial and basomedial amygdala and adjacent basal forebrain. Because correlation-based measures are sensitive to sample size, we interpret the magnitude of these effects descriptively; the robust observation is that dependence restructures the coordination of brain-wide activity without dismantling its global architecture. These two observations describe different levels of the network. The summary metrics index how much clustering and integration the network has, while hub identity indexes which regions occupy its most connected positions. A network can preserve the former while redistributing the latter, and that is what these data show: the global metrics are statistically indistinguishable within the precision this design allows, whereas hub membership changes reliably, with 24 of the 30 regions that lose hub status under dependence appearing as hubs in at least 10 of the 12 leave-one-animal-out vehicle networks and in no more than 3 of the 10 dependent ones.

Several of these hub regions, notably the lateral hypothalamus and amygdala, also appear in the durable morphine-associated networks identified by resting-state fMRI in a mouse model of morphine abstinence (Sourty et al. 2024), although the strongest changes in that study centered on the retrosplenial cortex and default mode network. Because co-activation quantified across animals is related but not identical to within-subject functional connectivity these fMRI studies report, and because such small, deep nuclei are difficult to resolve with human imaging, whole-brain cellular mapping and fMRI are best regarded as complementary levels of description. Consistent with its weaker regional activation, Ro 64-6198 left interregional co-activation close to vehicle levels, reinforcing that NOP receptor activation produces a smaller change in both regional activation and interregional co-activation than morphine at the doses used.

Several features of the design constrain interpretation. Vehicle animals received a single injection and are therefore injection-matched to the acute morphine and Ro 64-6198 conditions, but not to the morphine-dependent condition, which involved repeated handling and injection across five days. Therefore, the dependent-versus-vehicle comparison confounds the effects of chronic morphine with those of repeated injection, and because injection-evoked Fos both adds activation and habituates with repetition, the direction of this confound cannot be determined a priori. This issue was somewhat diminished by prior handling with four i.p. injections of saline to vehicle and drug-treated mice. The dependent condition also differs from the acute condition in dose as well as in exposure history, which was intentional and necessary to interrogate the two conditions. Estrous cycle was not tracked, and 4-hydroxytamoxifen is itself a pharmacological agent with affinity for estrogen receptors; the female regional signature reported here lies in estrogen-sensitive nuclei, and an interaction between 4-OHT and hormonal state at these sites cannot be excluded as a contributor. Changes in locomotion were observed but not recorded, because mice were confined within the testing chambers to reduce further neuronal activation. In rodents, opioids increase locomotor activity whereas NOP agonists tend to reduce it, so motor-related activation may contribute to differences in global neuronal activity between conditions and cannot be fully separated from other drug effects. The analyzed set also excludes the ventral-most structures, where the mounting adhesive impedes light transmission. Among them is the rostral ventromedial medulla, so the descending pain-modulatory pathway is represented here by the periaqueductal grey, pedunculopontine and pontine reticular nuclei but not by its medullary output stage, and all regional and network results describe the 198 analyzed regions rather than the whole grey-matter brain (Table S1; Supplementary Methods). Finally, the morphine and Ro 64-6198 doses were matched on published antinociceptive data rather than by behavioral testing in these animals, which would itself have driven c-Fos expression.

Removing each animal’s global activation level before computing correlations — global signal regression, by analogy with functional-connectivity MRI (Murphy and Fox 2017) — is the most aggressive correction these data permit, and every co-activation result is reported both with and without it (Figure S1d,e). We retain the uncorrected version as primary because the global signal here is not a nuisance term: per-animal global activation itself differed across conditions (η² = 0.30), in the same pharmacological order as the regional effects, so approximately 30% of the variance in the quantity such a correction discards is itself attributable to condition, and regressing it out removes drug effect along with animal-level variance. This objection is long established in the imaging literature, where global signal regression is known to discard neurally meaningful variance alongside artifact (Liu et al. 2017; Murphy and Fox 2017). Read in the most conservative manner, what survives the correction is the redistribution of which region pairs co-vary under dependence, while the uniform increase in co-activation strength, and the greater regional breadth of the male response, do not.

Taken together, these results show that morphine, and to a lesser extent NOP receptor activation, alters brain-wide activity at three levels; the number and identity of activated regions, the strength of their coordination, and the composition of network hubs. The magnitude of the response differs by sex with canonical reward circuitry (NAc, VTA, BNST) and midbrain reaching nominal significance in males but not in females at this dose; however, these may reflect differences in detection as well as sex specificity. The convergence of the regional and network analyses on hypothalamic and amygdalar circuitry, and the emergence of heightened interregional co-activation as the dominant signature of dependence, mark these features as priorities for mechanistic follow-up. More broadly, this work establishes whole-brain, cellular-resolution activation mapping as a framework for dissecting opioid action across the entire brain. Set against the weaker, modulatory signature of NOP receptor activation, these results begin to define how distinct members of the opioid receptor family alter brain-wide activation, a distinction that will grow in importance as bifunctional NOP/mu compounds advance toward the clinic (Kiguchi et al. 2022; Jansen et al. 2026). Because the TRAP2/Ai9 strategy permanently tags the neurons engaged under each condition, the same strategy applied without terminal imaging would render these populations accessible to causal, circuit-level manipulation, and the maps presented here provide a whole-brain, cellular-resolution foundation for resolving how opioids and NOP ligands remodel brain activity across analgesia, tolerance, and dependence.

## Supporting information

Table S5 (ESM_6)

Table S4 (ESM_5)

Table S3 (ESM_4)

Table S2 (ESM_3)

Tale S1 (ESM_2)

## Declarations

### Ethics approval

All animal procedures were approved by the Florida Atlantic University Institutional Animal Care and Use Committee (protocol no. A24-06) and conducted in accordance with the NIH Guide for the Care and Use of Laboratory Animals; the study is reported in accordance with the ARRIVE guidelines.

### Data availability

All processed data needed to reproduce the analyses and figures, including the per-region cell-count and density matrices and the statistical and network result workbooks, are deposited in Zenodo (https://doi.org/10.5281/zenodo.23811912; Martinez et al. 2026c). All analysis code (R) is maintained in a public GitHub repository (https://github.com/toll-lab-code/cfos-opioid-brain-analyses) and archived in Zenodo (https://doi.org/10.5281/zenodo.23811901; Martinez et al. 2026a). The dataset and code are also cited in the reference list. Raw light-sheet imaging volumes are not deposited in a public repository owing to their size (on the order of terabytes per brain); this size exception is noted here, and the volumes are available from the corresponding author on reasonable request. The Allen Mouse Brain Common Coordinate Framework (CCFv3) is publicly available, and cell detection and atlas registration were performed in NeuroInfo (MBF Bioscience).

### Authors’ contributions

L.T., M.M., and A.O. conceptualized the study. M.M., J.T., and A.O. performed the investigation. M.M. and D.V.Z. carried out the formal analysis. L.T. and A.O. acquired funding. M.M. and L.T. wrote the manuscript. All authors read and approved the final version of the manuscript.

## Figures and Supplementary Material

Supplementary material is provided as six Online Resources: the Supplementary Information document (Online Resource 1), which contains the Supplementary Methods, Supplementary Results and Supplementary Figures S1–S3, and Supplementary Tables S1–S5 as separate spreadsheets (Online Resources 2–6).

**Online Resource 1 (ESM_1.pdf)**

Supplementary Information: Supplementary Methods, Supplementary Results, and Supplementary Figures S1–S3.

**Online Resource 2 (ESM_2.xlsx)**

Table S1, region key. Abbreviation, full structure name and CCFv3 division for all 246 candidate grey-matter regions, with inclusion status (198 analyzed, 48 excluded), the reason for each exclusion in its two classes, and the per-structure registration coefficient of variation.

**Online Resource 3 (ESM_3.xlsx)**

Table S2, per-region results. One-way ANOVA on log10 c-Fos density for each of the 198 analyzed regions, with Cohen’s d, Hedges’ g, 95% confidence intervals, Tukey-adjusted p values (uncorrected across regions) and Benjamini–Hochberg q values for every condition pair, combined and by sex, together with the two-way condition × sex terms.

**Online Resource 4 (ESM_4.xlsx)**

Table S3, differential co-activation. The 176 region pairs whose co-activation differed significantly between conditions (Fisher r-to-z, Benjamini–Hochberg q < 0.05) across six comparison × sex groups, with both correlations, Δr, Fisher Z and q.

**Online Resource 5 (ESM_5.xlsx)**

Table S4, global network metrics. Mean absolute co-activation, edge threshold, mean node strength, weighted transitivity, global efficiency, modularity Q and Z, and small-world σ for each of the 12 condition × sex-group networks, with 95% bootstrap intervals on the combined rows.

**Online Resource 6 (ESM_6.xlsx)**

Table S5, per-region network roles. Guimerà–Amaral role, module identity and size, within-module degree z-score, participation coefficient and multi-centrality hub status for each of the 198 regions under each of the four conditions, combined and by sex.

