## Supplementary Information for "Whole Brain Cellular Activation After Mu and NOP Receptor Agonism: Shared Regional Targets with Divergent Network Consequences"

*Psychopharmacology*

Madeline Martinez, Akihiko Ozawa, Daniel Van Zant, Jake Thornberry, Lawrence Toll

**Supplementary Methods**

### Animals

TRAP2 (Fos2A-iCreER) and Ai9 (Rosa26-LSL-tdTomato) mice (The Jackson Laboratory, stocks #030323 and #007909) were crossed in-house to generate TRAP2/Ai9 reporter mice (Targowska-Duda et al. 2024). Male and female mice were used for all cohorts (n = 6 per sex per condition, except Morphine-Dependent, n = 5). Animals were 6–8 weeks of age at the start of the experiments, group-housed (3–5 per cage) under a 12 h light/dark cycle (lights on at 07:00) with food and water available ad libitum. All procedures were approved by the Florida Atlantic University IACUC and conducted in accordance with the NIH Guide for the Care and Use of Laboratory Animals.

### Activity-dependent labeling and drug treatments

Neuronal activation was visualized genetically using TRAP2/Ai9 mice, in which tamoxifen inducible CreERT2 is expressed under the control of c-fos promoter. Administration of 4-hydroxytamoxifen (4-OHT) enables Cre-mediated recombination at the Ai9 locus (Madisen et al. 2010), resulting in permanent tdTomato expression in neurons that were transcriptionally active within the labeling window (Corder et al. 2019; Targowska-Duda et al. 2024). Mice were assigned to one of four conditions: Vehicle, Acute Morphine, Morphine-Dependent, or the nociceptin/orphanin FQ (NOP) receptor agonist Ro 64-6198. Morphine sulfate and Ro 64-6198 (both from the NIDA Drug Supply Program) were dissolved in vehicle (5% DMSO, 0.5% hydroxypropyl cellulose). Animals received a single drug administration for Acute Morphine (10 mg/kg, i.p.), Ro 64-6198 (0.6 mg/kg, i.p.) and vehicle 30 min before 4-OHT injection. For Morphine-Dependent condition, animals received an escalating, twice-daily (a.m./p.m., approximately 8 h apart) morphine regimen over four days (Day 1, 10/15 mg/kg; Day 2, 20/30 mg/kg; Day 3, 50/60 mg/kg; Day 4, 70/75 mg/kg), followed on Day 5 by a single morning challenge dose (25 mg/kg, i.p.) 30 min before 4-OHT injection. The challenge dose was intended to limit withdrawal at the time of labeling; withdrawal signs were not scored and its absence was not independently confirmed. 4-OHT (50 mg/kg, i.p., in corn oil; Allen et al. 2017) was administered 30 min after the drug/vehicle injection, at the period of peak drug effect, and mice were left undisturbed in the testing environment for 2 h. All animals were perfused 10 days after 4-OHT injection to allow robust tdTomato expression.

The two morphine conditions differ in the dose administered during the labeling window as well as in treatment history. Acute-morphine animals were labeled following a single 10 mg/kg injection, whereas morphine-dependent animals were labeled following the 25 mg/kg Day 5 challenge, which was given to ensure that animals were not in withdrawal at the time of labeling. The design therefore does not separate treatment history from challenge dose, and no acute 25 mg/kg control group was included. Because dependent animals received the larger challenge dose yet showed lower activation than acutely treated animals across cerebellar, pontine and medullary structures, the reduction reported for those regions is a conservative estimate of the effect of chronic treatment; the same reasoning does not apply in the opposite direction, and regions showing comparable activation across the two conditions should not be read as evidence of preserved sensitivity.

**Tissue clearing and light-sheet imaging**

Whole brains were cleared (not immunolabeled) using MDISCO, a fluorescence-preserving solvent-based clearing protocol (Martinez et al. 2026), and imaged intact on a Miltenyi UltraMicroscope Blaze II light-sheet fluorescence microscope using uniform acquisition parameters across samples. Mice were transcardially perfused with 20–30 mL of 1× PBS followed by 20–30 mL of freshly prepared, filtered 4% (w/v) paraformaldehyde (PFA) in PBS (pH 7.4); brains were dissected, post-fixed in 4% PFA overnight at 4 °C, and washed in 1× PBS overnight at 37 °C. Fixed brains were dehydrated through a graded methanol (MeOH) series in water (20%, 40%, 60%, and 80% v/v, 1 h each at room temperature [RT]) followed by two changes of 100% MeOH (1 h, then overnight at RT). Samples were delipidated and permeabilized in 66% dichloromethane (DCM)/33% MeOH (v/v) overnight at RT, returned to 100% MeOH (two 2-h washes at RT), and bleached in freshly prepared 6% hydrogen peroxide in MeOH (one part 30% $H_2O_2$ to four parts MeOH) overnight at 4 °C to reduce background. After post-bleaching MeOH washes (1 h, then overnight at RT), brains underwent a final delipidation in 66% DCM/33% MeOH (3 h at RT) and 100% DCM (1 h at RT). For refractive-index (RI) matching, samples were transferred to 100% dibenzyl ether (DBE) and rocked at RT until optically transparent, held in DBE for 4–5 days to allow internal bubbles to dissipate, and finally transferred to ethyl cinnamate (ECi) for final RI matching, imaging, and long-term storage. These MDISCO-specific modifications preserve endogenous tdTomato fluorescence and yield uniform optical transparency without antibody-based signal amplification. Tiled acquisitions were stitched into whole-brain volumes with MACS iQ View (Miltenyi Biotec).

Stitched volumes were inspected in Imaris (Oxford Instruments) for reconstruction quality and signal uniformity and for preparation of representative images only (no quantitative measures were derived in Imaris).

**Atlas registration, region cleaning, and quantification**

Whole-brain volumes were processed in NeuroInfo (Tappan et al. 2019) for automated cell detection, registration, and region-based quantification. Volumes were registered to the Allen Mouse Brain Common Coordinate Framework (CCFv3) by an intensity-based approach (coarse manual alignment followed by nonlinear registration). tdTomato-positive somata were detected automatically using a size constraint of 10–35 µm diameter and assigned to CCFv3 structures; every brain's registration was visually inspected against the CCFv3 reference and confirmed before export, with detections reviewed in the same pass. Registration accuracy is not uniform across the brain: the validation reported by the developers of this software indicates that large and dorsally located regions are delineated most accurately (Tappan et al. 2019), so alignment of small ventral and midline structures carries correspondingly greater uncertainty. This is a general property of registering cleared whole-brain volumes to a histology-derived reference rather than a feature of this dataset, and interpretation of the smallest ventral and midline regions has been limited accordingly. Per-structure registration consistency is reported in Table S1 as the coefficient of variation of each structure's registration-derived volume across brains, taken from the validation published with the software (Tappan et al. 2019); the median across the 198 analyzed regions is 0.085. Among the regions on which the conclusions rest, the mammillary body is among the most consistently delineated (0.069), whereas the accessory supraoptic group (0.122) and the presubiculum (0.156) are the least, and those two are interpreted with that in mind. NeuroInfo also returns a per-brain alignment score; we do not report it because it has no published reference range against which a value could be interpreted.

NeuroInfo returns counts for all CCF structures, including white matter, fiber tracts, and ventricular spaces. The exported output was cleaned to the analyzed region set as follows: (i) only grey-matter regions were retained; (ii) left and right hemisphere counts were averaged to a single bilateral value per region; and (iii) the ventral-most regions, where the brain is adhered to the imaging stage and the mounting adhesive impedes light transmission and yields unreliable signal, were excluded. This yielded a final analyzed set of 198 grey-matter regions; the complete list of included and excluded regions is provided in Table S1. Region volumes used for density normalization were the mean registration-derived CCFv3 structure volumes reported by the developers of this software (Tappan et al. 2019), rather than the nominal atlas volumes (Wang et al. 2020). Across the analyzed set these are on average 15% smaller than the nominal volumes and therefore absorb the mean deformation introduced by clearing and registration. Because a region's volume is a constant across animals, this choice shifts absolute densities but leaves every between-condition comparison unchanged.

The a priori exclusions listed in Table S1 include several structures with established roles in opioid action. Among them are the nucleus raphe magnus, nucleus raphe pallidus and nucleus raphe obscurus, which together with adjacent reticular nuclei constitute the rostral ventromedial medulla; the arcuate, ventromedial and supraoptic hypothalamic nuclei; and the ventrolateral and ventromedial preoptic nuclei. These regions lie at or near the ventral surface, where the mounting adhesive impedes light transmission, and were excluded on that technical basis rather than on any anatomical or functional criterion. Regional and network results should accordingly be read as describing the 198 analyzed regions rather than the whole grey-matter brain. The descending pain-modulatory pathway in particular is represented in the analyzed set by the periaqueductal grey, pedunculopontine nucleus and pontine reticular nuclei, but not by the rostral ventromedial medulla itself, and the hypothalamic findings reported here are drawn from the 27 hypothalamic regions that survived exclusion rather than from the full hypothalamus.

Regions were assigned to anatomical divisions following the Allen Mouse Brain Common Coordinate Framework (CCFv3) ontology, with two functional groupings: amygdalar nuclei were treated as a single Amygdala division rather than divided between the cortical subplate and striatum as in the strict CCFv3 hierarchy, and the lateral septal complex was grouped with the pallidum (Pallidum/Septum). The amygdalar grouping reflects the functional coherence of the amygdalar complex, which is recruited as a unit in drug-related brain-wide co-activation (Kimbrough et al. 2020). It affects only the anatomical color-coding and the descriptive within-division co-activation summary (Figure 5d), not the regional statistical models or the network metrics, which operate at the region and module level. The brainstem was resolved into separate Pons, Medulla, and Cerebellum divisions, yielding 13 anatomical divisions across the 198 analyzed regions. For the force-directed (Kamada-Kawai; Kamada and Kawai 1989) network graphs (Figure 6), the 13 anatomical divisions were collapsed to seven major Allen divisions (Cerebral cortex: isocortex, olfactory, hippocampus, and cortical subplate; Cerebral nuclei: amygdala, striatum, and pallidum/septum; Thalamus; Hypothalamus; Midbrain; Hindbrain: pons and medulla; and Cerebellum) for legibility, since 13 node-fill colors are not reliably distinguishable in the dense graph layouts. All other panels, including the correlation heatmaps and the cartographic role plots, retain the full 13-division scheme.

### Normalization and regional statistical models

Per-region tdTomato+ neuron counts were exported separately for males and females. Counts were converted to density (cells/mm³) using CCFv3 region volumes (Wang et al. 2020), with zero values set to missing prior to $\log_{10}$ transformation. Because $\log_{10}(0)$ is undefined, animals with a zero count in a region were excluded from that region's ANOVA (1 value in males and 17 in females, across 11 regions), with the number of animals entering and excluded per region reported in the deposited workbooks. The primary regional test was a one-way ANOVA on $\log_{10}$ density with condition as the sole factor and males and females pooled, followed by Tukey-adjusted pairwise contrasts for the five a priori comparisons; p-values were corrected across the 198 regions by the Benjamini-Hochberg procedure (Benjamini and Hochberg 1995). The Morphine-Dependent versus Ro 64-6198 contrast, the sixth possible pairwise comparison, was excluded a priori because these conditions differ simultaneously in receptor target (mu versus NOP) and in treatment history (chronic versus acute), so it confounds the two factors and does not test an interpretable hypothesis. As secondary analyses, the one-way ANOVA was repeated within each sex (males and females analyzed separately). A condition × sex two-way ANOVA on $\log_{10}$ density was run as a supplemental analysis, testing the sex main effect and the condition × sex interaction per region (each Benjamini-Hochberg corrected across regions) alongside the condition contrasts. For each comparison we report Cohen's d (computed on the $\log_{10}$-density scale) and the Benjamini-Hochberg FDR q-value from the pooled, male-only, and female-only tests and from the negative-binomial GLM robustness model, together with the omnibus ANOVA F and its FDR q and the condition × sex (sex main-effect and interaction) q-values (Table S2). Benjamini-Hochberg FDR ($q < 0.05$) is the inferential criterion throughout; the sex-stratified ANOVAs are secondary/exploratory, and where FDR returned no significant regions within a sex-stratified panel, Tukey-adjusted p-values (not corrected across regions) are shown descriptively only — explicitly flagged as not surviving FDR correction — rather than as inferential claims. Counts of regions reaching Tukey $p < 0.05$ without surviving FDR were additionally calibrated against their own null, because the 198 regions are strongly correlated and such a count therefore cannot be read against the nominal 5% expectation. Condition labels were permuted within sex, preserving the sex composition of every group and the across-region correlation structure; the per-region ANOVA and Tukey contrast were recomputed on each permutation; and the observed count was placed in the resulting null distribution (10,000 permutations, add-one permutation p-value). Observed counts, null medians, 95th and 99th percentiles, null maxima and permutation p-values for every stratum and pairwise contrast are reported in the deposited workbook (Drug_Region_Count_Permutation.xlsx). Global activation was quantified per animal as the mean log10 regional density across the analyzed regions — the same per-animal global offset used in the co-activation controls — and compared across the four conditions by one-way ANOVA with Tukey post hoc contrasts, both pooled and within each sex, with a two-way condition × sex ANOVA reported alongside. The same analysis was repeated on the raw whole-brain tdTomato+ total and on a mean restricted to the regions labeled in every animal; all three are reported in the deposited workbook (Drug_Global_Activation.xlsx). The ordering of conditions was identical across measures, but the raw total, which is dominated by the largest structures, gave uniformly weaker contrasts (acute morphine $p = 0.056$, morphine-dependent $p = 0.31$), so the morphine-dependent result is the less robust of the two. Cross-condition similarity of the regional effect profiles was quantified by correlating, across the 198 analyzed regions, the per-region Cohen's d values from each vehicle-referenced contrast (vehicle versus acute morphine, versus morphine-dependent, and versus Ro 64-6198). This asks whether two conditions engage the same regions in the same rank order, which counts of significant regions cannot address. Each profile was also regressed on the acute-morphine profile, the slope giving the amplitude ratio and the ratio of standard deviations indicating whether a profile is compressed or more dispersed than the acute-morphine profile; the number of regions falling below vehicle is reported alongside, because it distinguishes a uniformly scaled response from a bidirectional one. Because all three contrasts share the same vehicle group, sampling noise in the vehicle means is common to every profile and inflates all pairwise correlations by a similar amount; these correlations are therefore comparable with one another but are not unbiased estimates of profile similarity in absolute terms, and the interpretable quantity is their ordering. Pearson and Spearman coefficients, confidence intervals, regression slopes and the per-region profile values are reported in the deposited workbook (Drug_Profile_Similarity.xlsx).

Model assumptions were assessed per region: residuals departed from normality (Shapiro-Wilk $p < 0.05$) in 63 of 198 regions, whereas variances were homogeneous in all but 8 (Brown-Forsythe $p < 0.05$). As a quality-control check on the primary log-density ANOVA, a negative binomial GLM was fit per region on bilateral summed counts (hemisphere averages × 2; MASS::glm.nb) with a condition × sex design and log(region volume) as an offset, using a likelihood-ratio omnibus test and Tukey-adjusted emmeans contrasts on the condition factor. This count-based model does not assume log-normally distributed densities and accommodates overdispersion; it is reported as a robustness check rather than the primary analysis and was used to confirm that the direction and significance of the primary findings are not driven by the distributional assumptions of the ANOVA.

**Network construction**

For each condition × sex, region × animal matrices were cleaned by setting negative values to missing and imputing zeros with half the minimum nonzero value for that region (a commonly used substitution for log transformation of count/omics data; Wei et al. 2018), followed by $\log_{10}$ transformation. This differs from the regional analysis, where zeros are excluded rather than imputed, because network construction requires a complete value for every region in every animal to compute pairwise correlations, whereas a per-region ANOVA can simply omit the missing observation. Regions with no variance or no valid data were dropped. Interregional co-activation was quantified as the Pearson correlation between regions across animals. Networks were built from log10 regional cell counts, whereas the regional analyses use log10 cell density (counts divided by CCFv3 region volume). The distinction does not affect any network result: within a region the volume is a constant, so the two quantities differ by a fixed per-region offset on the log scale, and a correlation computed across animals is invariant to it. We verified this directly — adding an arbitrary per-region offset to the count matrices, log10 region volume being one such offset, leaves the correlation matrices unchanged to numerical precision (maximum absolute difference $2.3 \times 10^{-15}$ across all 19,503 region pairs, with no pair undefined; checked on every run, Scale_invariance sheet of 04_Coactivation_Controls.xlsx).

The primary analysis used a pooled Combined network. Because the per-sex sample size (n = 5–6) is too small to estimate stable interregional correlations, males and females were pooled after mean-centering each region within sex (Alexander-Bloch et al. 2013), which removes the between-sex mean shift that would otherwise induce spurious cross-region correlations while preserving the within-sex covariance the network relies on, and provides the sample size (n = 10–12) needed for stable network estimation and for bootstrap and permutation inference. Sex-stratified (Males, Females) networks were computed as a supplemental sensitivity analysis (Tables S3–S5); consistent with their limited power at this sample size they yielded few differentially co-activated pairs, and sex differences are examined at the regional level (Figure 4) rather than the network level.

**Thresholding**

Networks were proportionally (density-matched) thresholded by retaining the strongest positive correlations until a fixed proportion of possible edges survived, set to a common density of 10% across all conditions so that degree-, modularity-, and role-based metrics are compared at matched density rather than confounded by condition differences in overall correlation strength (Rubinov and Sporns 2010; van den Heuvel et al. 2017). Thresholding fixed the edge count identically across networks (1,951 edges); regions left with no suprathreshold edge were then excluded, so density among the retained regions ranged from 0.101 to 0.121 rather than a uniform 0.10. Only positive edges were retained (Chen et al. 2011; Kimbrough et al. 2021). The heatmaps in Figure 5a display the full correlation matrix, including negative values, whereas all graph metrics are computed on the positive-edge thresholded network. This proportional scheme is a departure from the fixed absolute-r threshold common in the c-Fos co-activation literature and is adopted to ensure cross-condition comparability of the graph metrics; mean absolute correlation (|r|) per condition is reported separately so that the proportional-thresholding caveat can be assessed.

**Module detection**

Communities were detected directly on the thresholded weighted graph by consensus Louvain (1,000 weighted-Louvain runs; co-classification matrix thresholded at 0.5 and re-clustered to convergence; Lancichinetti and Fortunato 2012), which addresses Louvain stochasticity and the modularity resolution limit and provides a stable, graph-native partition for the cartographic metrics. To document the basis for this choice, we also computed module counts using the standard c-Fos approach (Euclidean distance between regions' interregional-correlation profiles, complete-linkage hierarchical clustering, dendrogram cut at 0.3–0.7 of maximum tree height; Kimbrough et al. 2020; Ardinger et al. 2024) on the full correlation matrix; because a single tree cut depends on the chosen cut height and is subject to the modularity resolution limit, these counts were computed only as a comparison with the standard approach rather than as the basis for the cartographic analysis, for which the consensus-Louvain partition was used.

**Node roles and hubs**

Cartographic node roles were assigned from the within-module degree z-score (WMDz) and participation coefficient (Guimerà and Nunes Amaral 2005), computed on the (thresholded graph, consensus-module) pair. Roles were driven by the weighted WMDz (a unit-SD within-module z-score on this data), with the hub/non-hub boundary set at WMDz = 1.5, the lower bound of the canonical high-WMDz range (Cohen and D'Esposito 2016). We note that the value of 0.8 reported in the c-Fos lineage was derived from a compressed WMDz distribution and is not comparable to a unit-SD z-score; on these data 0.8 over-calls hubs (~25% of nodes), whereas 1.5 flags roughly 4% of nodes (~8 hubs per network). Both binary and weighted WMDz/participation variants are reported; weighted variants drive the role assignment. In this scheme, hubs are further distinguished by participation coefficient into provincial hubs (edges concentrated within their home module) and connector hubs (edges distributed across modules); the between-module connector hubs are the regions labeled in each network in Figure 6c, with the multi-centrality convergence below reported as independent supporting evidence. Roles for regions in modules smaller than five nodes are flagged as lower-reliability (small-module instability and participation-coefficient size bias; Pedersen et al. 2020).

Hub stability was assessed by leave-one-animal-out jackknife on the Combined networks. For each condition every animal was removed in turn, the correlation matrix rebuilt, thresholded at the same 10% density, and the multi-centrality hub set recomputed. Agreement with the full-sample hub set is summarized by the Jaccard index and by the proportion of jackknife networks in which each full-sample hub was recovered.

As convergent evidence independent of the modular partition, hubs were also identified by agreement across four centrality measures (strength, weighted betweenness, weighted closeness, and eigenvector centrality; Sporns et al. 2007; van den Heuvel and Sporns 2013). A region was scored "high" on a measure if it exceeded the within-network mean by more than one standard deviation and was classified a hub if high on at least two of the four measures. Eigenvector centrality was computed on the giant connected component, with off-component nodes set to missing; because the sex-stratified networks fragment into multiple components at matched density, this guard was necessary. A continuous hub-ness score (mean within-network z across the four centralities) is also reported for ranking.

**Global metrics and modularity significance**

Global metrics (mean strength, weighted transitivity, global efficiency, and Louvain modularity Q) were computed on the density-matched graph after isolate removal, with the number of components and giant-component fraction reported. Modularity significance was tested against a degree-preserving (configuration-model) null in which the observed graph was rewired preserving each node's degree (Maslov-Sneppen edge swaps; Rubinov and Sporns

2010); we report observed Q, the null mean and SD, a modularity z-score, and a permutation p-value (5,000 permutations). A small-world index (Humphries and Gurney 2008) was computed against 100 randomly rewired nulls (each-edge rewiring, which does not preserve the degree sequence). Both comparisons benchmark against rewired nulls that do not preserve the transitivity induced by correlation. Correlation-based networks are inherently more clustered than degree-matched random graphs for this reason, and degree-preserving rewiring in particular overestimates small-worldness (Zalesky et al. 2012). The modularity z-scores and small-world indices reported here are therefore interpreted as confirming that the networks are well formed rather than as evidence of biological organization, and no condition comparison rests on them.

**Co-activation differences**

Edge-wise condition differences in interregional co-activation were tested by Fisher r-to-z transformation (two-sided), with Benjamini-Hochberg correction across all region pairs, for the same five a priori condition contrasts; the per-pair correlation difference (Δr) is reported alongside. Co-activation structure is displayed per condition as full region × region correlation heatmaps, with regions ordered by anatomical division and decoded by a division color key (Figure 5a). Overall co-activation strength is summarized as the mean absolute correlation (|r|) per condition (Figure 5b), and within-division co-activation as the mean |r| among within-division region pairs for each division across the four conditions (Figure 5d); anatomical divisions with fewer than three regions (the cortical subplate) are omitted from the within-division summary. The number of region pairs strengthened under dependence is ranked by region (Figure 5c). These heatmaps and division summaries are descriptive; inferential testing of condition differences in co-activation is provided by the edge-wise Fisher r-to-z analysis above.

**Sensitivity of co-activation strength to animal-level variance**

Because interregional co-activation is computed across animals, any component of variance shared by all regions within an animal — for example, animal-to-animal differences in overall labeling efficiency — contributes to every region pair and therefore raises the mean absolute correlation independently of regional coordination. We quantified this by decomposing each condition's region × animal matrix into a per-animal global offset (the mean $\log_{10}$ density across regions) and the region-specific residual. The proportion of variance attributable to the global offset was 29.0% under vehicle, 41.7% under acute morphine, 62.6% under morphine dependence and 33.9% under Ro 64-6198. The elevated value in dependent animals reflects reduced region-specific variance (SD 0.123, versus 0.266 under vehicle) rather than increased global variance, which was comparable across conditions (SD 0.167 versus 0.178; Variance_decomposition sheet of the deposited workbook 04_Coactivation_Controls.xlsx). Removing the per-animal offset before computing correlations — global signal regression (GSR), the analogue in this design of the procedure used in functional-connectivity MRI, with animals in place of timepoints and regions in place of voxels (Liu et al. 2017; Murphy and Fox 2017) — reduced mean $|r|$ to 0.390 (vehicle), 0.310 (acute morphine), 0.376 (morphine-dependent) and 0.354 (Ro 64-6198); median-centring gave equivalent results. The increase in co-activation under dependence therefore reflects a shift in the ratio of animal-level to region-specific variance — that is, activation profiles that are more stereotyped across dependent animals — as much as an increase in coupling between specific region pairs, and the two cannot be separated at this sample size. Values without GSR are reported throughout for comparability with the c-Fos co-activation literature, in which per-animal normalization is not applied (Kimbrough et al. 2020; Ardinger et al. 2024), and the dependence effect is interpreted accordingly.

Two further controls were applied to the co-activation analysis. First, because correlation magnitude depends on sample size and the morphine-dependent group was smaller (n = 5 per sex) than the others (n = 6 per sex), all conditions were subsampled to 5 animals per sex, the network rebuilt, and the mean absolute correlation recomputed over 2,000 random draws, summarized as the mean and the 2.5th–97.5th percentiles. Second, the vehicle versus morphine-dependent difference in mean |r| was tested by label permutation, pooling animals within sex, reassigning condition labels at the observed group sizes, and recomputing the difference under the null (2,000 permutations, two-sided). Both controls were run on the uncorrected matrices and repeated after removal of the per-animal global offset.

**Inference on network metrics**

Because each condition yields a single correlation network, global metrics are point estimates without intrinsic error bars. To support inferential claims, we resampled over animals (the unit of replication) on the Combined networks: (i) bootstrap confidence intervals were obtained by resampling animals with replacement, rebuilding the network at matched density, and recomputing each metric (1,000 iterations), yielding a 95% CI per condition; and (ii) between-condition differences were tested by label-permutation, pooling animals, shuffling condition labels at the observed group sizes, and recomputing the metric difference under the null (two-sided p; 1,000 permutations; Vetere et al. 2017; Kimbrough et al. 2020). With small samples, bootstrap resampling can duplicate animals and produce wide intervals; this width reflects the sample size honestly. Benjamini-Hochberg correction was applied within two metric families analyzed separately: a Strength family (mean strength, which re-expresses co-activation strength at matched density) and a Topology family (weighted transitivity, global efficiency, modularity Q), mirroring the lineage's separation of correlation strength from graph topology (Kimbrough et al. 2020; Bloch et al. 2021).

**Software and packages**

Analyses were performed in R 4.4.3. Regional models used readxl, openxlsx, dplyr, MASS, and emmeans; network analyses used tidyverse and igraph. Figures used ggplot2 with showtext, ggbeeswarm, and scales (regional figures); base R graphics (network co-activation heatmaps, division summaries, and force-directed network graphs); and jsonlite, xml2, rsvg, magick, and viridisLite (atlas projections). Coronal section geometry for the atlas projections (Figure 6a,b) is from the Scalable Brain Atlas template for the Allen Mouse Brain Common Coordinate Framework v3 (ABA_v3; Lein et al. 2007; Bakker et al. 2015); participation coefficient and WMDz were matched to atlas regions by acronym, and regions without a value in a given condition are shown in grey. All stochastic procedures (consensus Louvain, modularity permutation, the region-count permutation, bootstrap and label-permutation resampling, and the force-directed graph layout) were run under fixed random seeds, so the deposited code reproduces the reported values exactly. Results were exported as topic-grouped Excel workbooks (00_Processed_Data, 01_Connectivity, 02_Network_Metrics, 03_Modules_and_Roles, 04_Coactivation_Controls), the regional statistical workbooks (Drug_Statistical_Results_Primary, BySex, TwoWay, NegBin, GlobalNorm and GlobalNorm_BySex), and Drug_Global_Activation, Drug_GlobalNorm_Concordance, Drug_Region_Count_Permutation, Drug_Profile_Similarity and Model_Assumption_Checks.

**AI use statement**

Generative AI (Claude, Anthropic; model Claude Opus 5.0) was used during manuscript preparation and in preparing the analysis code for public deposit. Specifically, it was used to: (i) correct, update, and debug analysis code written by the authors, and to write utility scripts used to assemble the supplemental tables; (ii) consistency-check manuscript and supplemental text, including terminology standardization and cross-checking reported numerical values against the source data workbooks; and (iii) prepare the deposited R code for public release, including removing machine-specific file paths, generalizing input handling, standardizing comments and headers, and drafting repository documentation. The study design, the analytical approach, the execution of all statistical and network analyses, and the interpretation of results were determined and carried out by the authors. No generative AI image tools were used to create figure content; all figure content was rendered directly from the study data by the analysis code. All AI-assisted outputs were reviewed, verified, and where necessary corrected by the authors, who take full responsibility for the accuracy and integrity of the work.

### Supplementary Results

#### Robustness of the regional analysis

Two sensitivity models were fitted alongside the primary $\log_{10}$-density ANOVA. The first, a negative binomial GLM on bilateral summed counts with log(region volume) as an offset, relaxes the log-normality assumption of the primary model. The second re-ran the one-way ANOVA on globally normalized densities, in which each region is expressed relative to that animal's brain-wide activation, and therefore asks whether a region is engaged disproportionately rather than in absolute terms. Each is more permissive than the primary analysis for some contrasts and more conservative for others. Both are reported here as sensitivity analyses; Benjamini–Hochberg-corrected inference throughout the manuscript is based on the primary ANOVA, which is the more conservative model. The two global corrections used in this manuscript are deliberately different. Here each region is expressed relative to that animal's whole-brain total, as a compositional robustness check on the regional models. In the co-activation analysis the per-animal global offset is instead removed by subtraction (global signal regression; see the co-activation controls above). A term shared by all regions within an animal inflates every pairwise correlation irrespective of whether those regions are coordinated, and is therefore a nuisance for the network analysis; for a between-condition comparison within a single region the same term is absorbed into the group means and is itself part of the treatment effect. No correction of either kind is applied to the primary regional analysis.

Both models reproduced the principal regional findings. The negative binomial GLM returned more FDR-significant regions than the primary ANOVA for the vehicle contrasts (93 versus 44 under acute morphine; 35 versus 15 under dependence, and one region versus none under Ro 64-6198), replicating 43 of the 44 acute-morphine regions with 99.5% sign agreement across all 198 regions. Global normalization, which by construction removes any brain-wide change in activation, left only the accessory supraoptic group and the mammillary body FDR-significant under acute morphine, both of them among the 44, consistent with the acute-morphine increase being predominantly global rather than regionally selective. Under dependence, by contrast, 12 regions remained FDR-significant after global normalization, 11 of them among the 15 identified by the primary analysis, indicating that the periventricular–preoptic core is engaged disproportionately and not merely as part of a brain-wide rise.

The same normalization applied within sex indicates that the sex difference in regional breadth is likewise global in origin. Under acute morphine, none of the 20 regions reaching FDR significance in males survived global normalization, and no region reached FDR significance in either sex. At the uncorrected Tukey threshold the direction of the difference reversed: male detections fell from 82 regions to 13 while female detections rose from 9 to 24, indicating that the greater breadth of the male response in absolute terms follows from the larger male global increase rather than from a wider set of regions being engaged. Consistent with this, the globally normalized per-region effects of the two sexes are positively correlated across all 198 regions ($r = 0.39$ under acute morphine and $r = 0.58$ under dependence, both $p < 0.001$, on the $\log_{10}$ mean-difference scale; $r = 0.13$ and $r = 0.48$ respectively when computed on Cohen's d, which divides each regional effect by a within-sex standard deviation estimated from five to six animals and so carries more noise). Correlations for the absolute-density analysis and for every pairwise contrast are reported in the deposited workbook (Drug_GlobalNorm_Concordance.xlsx). Because each sex's per-region effect rests on five to six animals, these correlations are attenuated by sampling noise and are lower bounds on the agreement between sexes; no correction for measurement error is applied, because the reliability of a per-sex effect vector cannot be estimated stably from five to six animals and such a correction would roughly double the reported correlation on the strength of that estimate. Together with the absence of any condition × sex interaction surviving correction, this indicates a shared regional distribution that differs in magnitude rather than in location.

The two sensitivity models converge on a single point of difference from the primary analysis. For the acute morphine versus morphine-dependent contrast, where the primary ANOVA identified no region surviving FDR correction, the negative binomial GLM identified seven regions (COPY, PRM, PYR, UVU, AT, NLL and ECU) and global normalization identified four (COPY, PRM, UVU and AT, a subset of the seven), in every case with lower activation in dependent animals and in every case a cerebellar, pontine or tegmental structure. These are the same regions described above as showing lower c-Fos+ density in dependent animals at Tukey $p < 0.05$. Because the primary $\log_{10}$-density ANOVA is the more conservative model, this reduction is reported throughout as a non-significant trend; that two independently specified sensitivity models identify it nonetheless suggests it is unlikely to be noise, and it is flagged here for that reason. The regional effect profiles were also compared directly across conditions. The Ro 64-6198 profile correlated with the acute-morphine profile at $r = 0.63$ (95% CI 0.53 to 0.70; Spearman rho = 0.65) across the 198 regions, more strongly than the dependence profile did ($r = 0.35$, CI 0.22 to 0.47; rho = 0.26), and its slope on the acute-morphine profile was 0.40 with 0.64 times the dispersion. Ro 64-6198 therefore reproduces the acute-morphine regional pattern at reduced amplitude, falling below vehicle in only 1 of 198 regions, whereas dependence departs from it in shape as well as in amplitude, falling below vehicle in 21 regions (10.6%) with 1.36 times the dispersion. That the rank correlation slightly exceeds the product-moment correlation for Ro 64-6198 indicates the similarity is not driven by a few extreme regions.

**Sensitivity of the co-activation analysis to sample size**

Because the morphine-dependent group contributed fewer animals than the other conditions, the elevated co-activation observed under dependence could in principle reflect sample size rather than treatment. Subsampling every condition to the dependent-group n (5 per sex, 2,000 draws) excluded this explanation. Dependent-group co-activation (mean $|r| = 0.74$) exceeded the 97.5th percentile of every subsampled control distribution: vehicle 0.47 (95% range 0.41–0.53), acute morphine 0.52 (0.42–0.57) and Ro 64-6198 0.51 (0.41–0.59). A label-permutation test on the vehicle versus dependence difference gave $p = 0.002$ (2,000 permutations). Matching group n for the correlation calculations follows the approach taken in the alcohol-dependence c-Fos literature (Kimbrough et al. 2020).

Removing the per-animal global offset attenuated the uniform component of the dependence effect but not the edge-level differences. Mean |r| under dependence fell to 0.38, comparable to vehicle (0.39), and the permutation test on that difference was no longer significant ($p = 0.90$). The edge-wise comparison nonetheless identified a similar number of altered region pairs (136 at $q < 0.05$, versus 160 in the uncorrected analysis), but their direction was no longer uniform: 68 pairs were more strongly and 68 less strongly co-activated under dependence, against 160 and none respectively in the uncorrected analysis (Figure S1e). Dependence therefore redistributes interregional co-activation, while the increase in overall co-activation strength additionally reflects the more stereotyped whole-brain activation profiles described in Supplementary Methods. The same analysis applied within sex accounts for the apparent sex difference in dependence co-activation. Mean |r| under dependence was higher in females than in males (0.833 versus 0.616), but after removal of the per-animal global offset the two were indistinguishable (0.486 versus 0.481), indicating that the larger female value reflects animal-level variation in overall activation rather than greater region-specific coordination. Per-sex values under both pipelines are shown in Figure S1d.

**Stability of hub assignment**

Hub assignment was stable at the level of both the hub set and the specific claims made. Across conditions the jackknife hub sets agreed with the full-sample set with mean Jaccard indices of 0.82 (vehicle), 0.74 (acute morphine), 0.66 (morphine-dependent) and 0.56 (Ro 64-6198), with mean recovery of individual full-sample hubs of 89%, 86%, 80% and 75% respectively. Prelimbic cortex was recovered in 92–100% of jackknife networks in every condition and anterior cingulate in 50–100%; the lowest value is anterior cingulate under dependence, recovered in only half of the dependent jackknife networks, and its hub status in that condition should be regarded as unstable. The regions that lost multi-centrality hub status under dependence did so consistently: nucleus accumbens (striatum), anterior olfactory nucleus, claustrum, ectorhinal, entorhinal and auditory areas and the bed nucleus of the stria terminalis were hubs in 83–100% of vehicle jackknife networks and in 0–30% of dependent ones. The regions that gained hub status under dependence include the lateral hypothalamic area, perifornical nucleus, zona incerta and posterior amygdalar nucleus, each recovered in every dependent jackknife network. Hub assignment under Ro 64-6198 is the least stable of the four conditions and those hub claims are interpreted accordingly. Nucleus accumbens is nonetheless a cartographic connector hub under dependence (Figure 6c). The two criteria are distinct - the multi-centrality criterion requires agreement across four centrality measures, whereas the cartographic criterion requires a high within-module degree z-score with connections distributed across modules - and a region can satisfy either without the other.

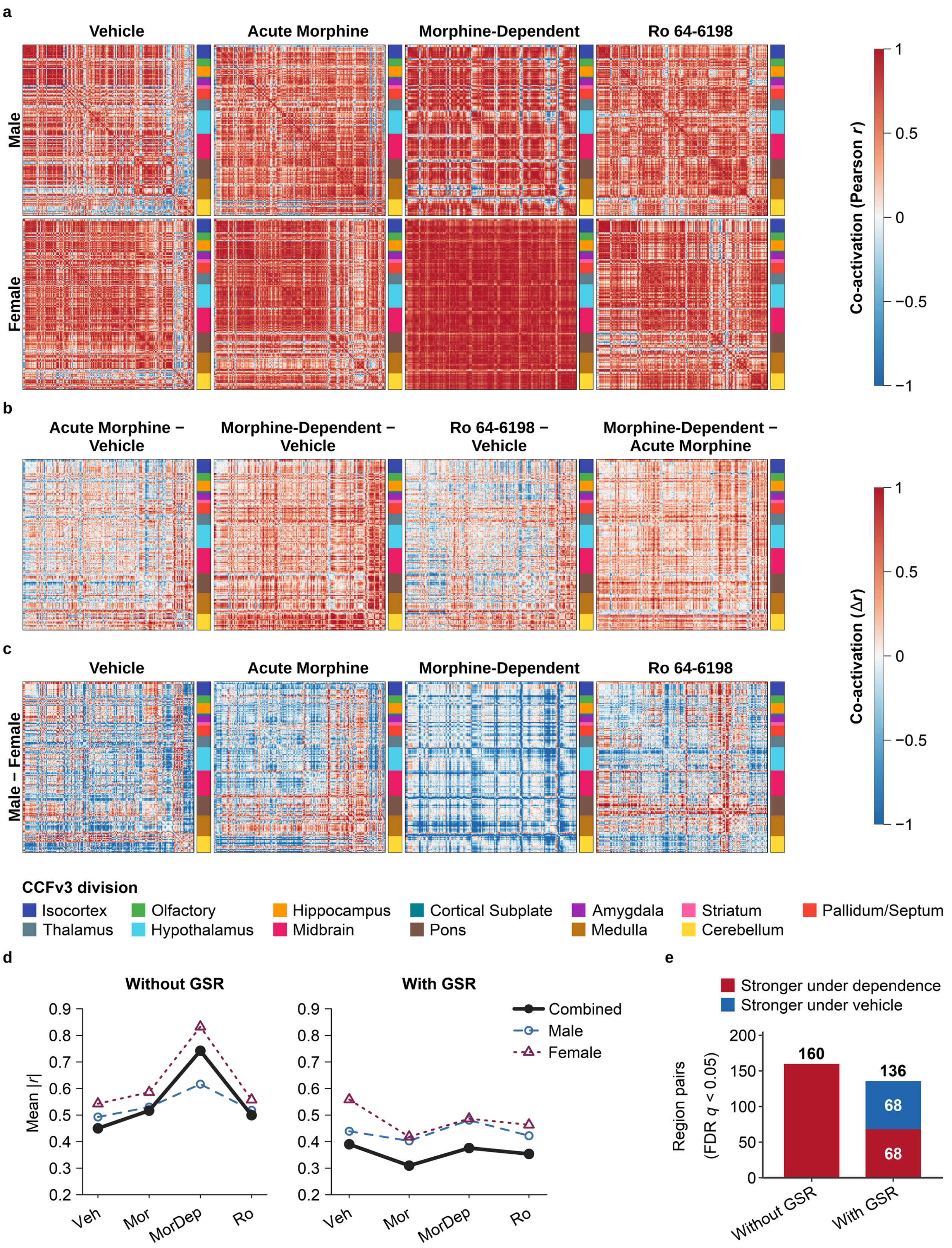
a
Vehicle
Acute Morphine
Morphine-Dependent
Ro 64-6198
Male
Female
Co-activation (Pearson r)
1
0.5
0
−0.5
−1
b
Acute Morphine − Vehicle
Morphine-Dependent − Vehicle
Ro 64-6198 − Vehicle
Morphine-Dependent − Acute Morphine
Co-activation (Δr)
c
Vehicle
Acute Morphine
Morphine-Dependent
Ro 64-6198
Male − Female
CCFv3 division
Isocortex
Olfactory
Hippocampus
Cortical Subplate
Amygdala
Striatum
Pallidum/Septum
Thalamus
Hypothalamus
Midbrain
Pons
Medulla
Cerebellum
d
Without GSR
With GSR
Mean |r|
Veh
Mor
MorDep
Ro
Combined
Male
Female
e
Stronger under dependence
Stronger under vehicle
Region pairs (FDR q < 0.05)
160
136
68
68
Without GSR
With GSR

**Figure S1. Sex-stratified whole-brain co-activation and its modulation across opioid and NOP-receptor conditions, and the sensitivity of co-activation to animal-level global activation. Co-activation was quantified as the pairwise Pearson correlation of log10-transformed regional c-Fos$^+$ cell counts across animals, over the 198 grey-matter regions. In every matrix, regions are ordered by anatomical division; the 13 major CCFv3 divisions are marked by the color band beside each matrix and defined in the CCFv3 division key. (a) Co-activation matrices for each condition, shown separately for males (top row) and females (bottom row); color encodes Pearson r from −1 to 1 (scale, top right). (b) Difference matrices (Δr) of the combined matrices (sexes pooled after within-sex mean-centering), in the treatment-minus-comparator direction: Acute Morphine − Vehicle, Morphine-Dependent − Vehicle, Ro 64-6198 − Vehicle, and Morphine-Dependent − Acute Morphine; red indicates stronger co-activation in the first-named condition, blue in the comparator (Δr scale, right of b and c). The sign convention matches Figure 5 (treatment − vehicle, increase-positive). (c) Sex-difference matrices (male − female) for each condition; red indicates stronger co-activation in males, blue in females, on the same Δr scale as b. (d) Mean absolute interregional correlation (mean |r|) per condition without (left) and with (right) global signal regression (GSR), for the combined network (black) and for males (blue) and females (wine), plotted as in Figure 5b, whose combined series is the left-hand black line here; combined-sex n = 12/12/10/12. (e) Number of region pairs with significantly altered co-activation under morphine dependence relative to vehicle (Fisher r-to-z, Benjamini–Hochberg q < 0.05), without and with GSR, split by direction; red denotes pairs more strongly co-activated under dependence and blue those more strongly co-activated under vehicle, as on the Δr scale of Figure 5. Conditions are Vehicle, Acute Morphine (10 mg/kg), Morphine-Dependent (escalating twice-daily dosing with a day-5 challenge), and Ro 64-6198 (0.6 mg/kg, NOP agonist); n = 6 per sex per condition except Morphine-Dependent (n = 5 per sex).**

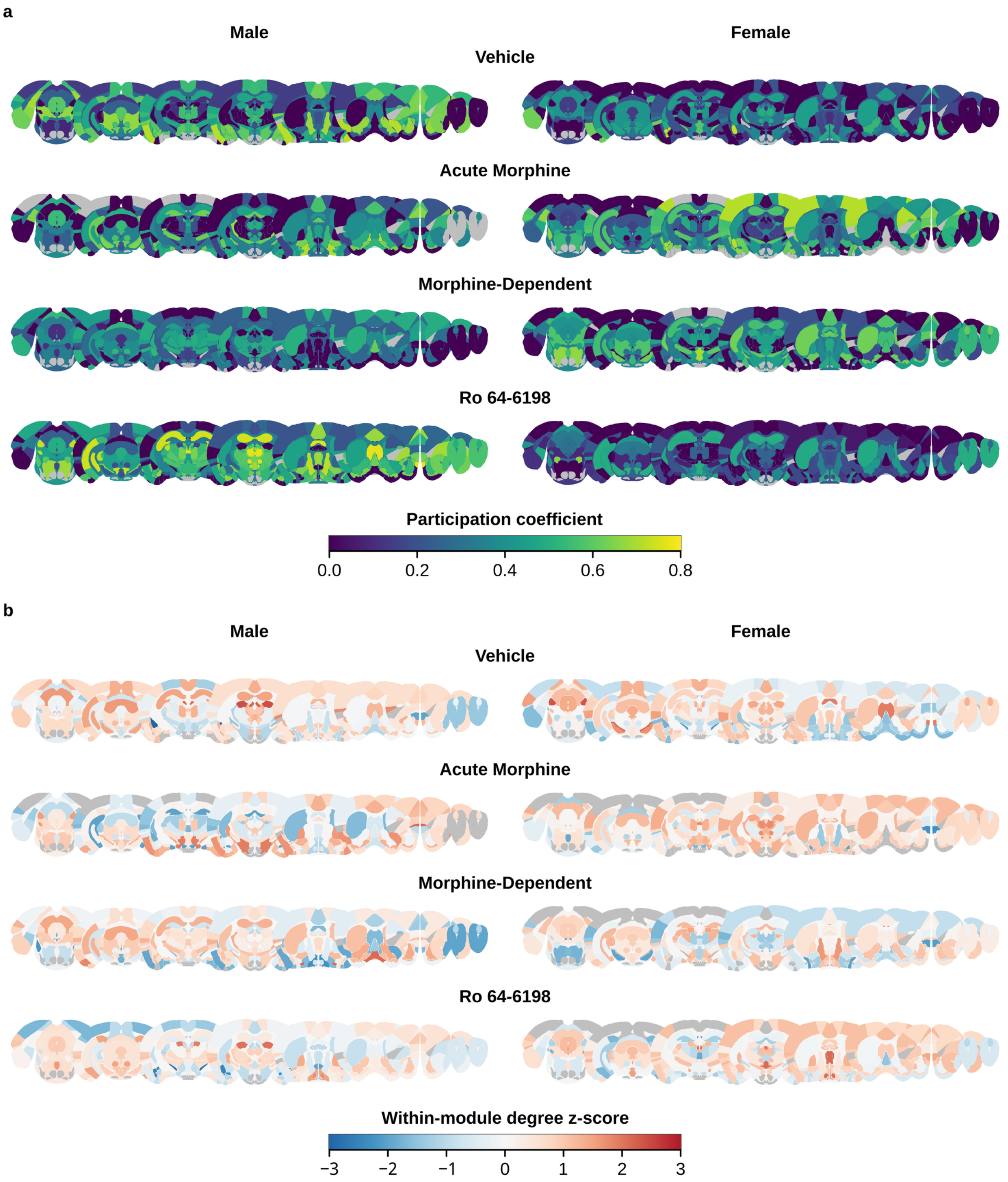


**Figure S2. Sex-stratified cartographic node metrics across the whole brain. Participation coefficient (a) and within-module degree z-score (WMDz) (b) for each condition (rows: Vehicle, Acute Morphine, Morphine-Dependent, Ro 64-6198), shown separately for males (left) and females (right). Metrics were computed on the sex-stratified co-activation networks (n = 6 per sex per condition except Morphine-Dependent, n = 5 per sex; 10% edge-density threshold; consensus-Louvain modules) and projected onto nine coronal sections spanning anterior to posterior, using Scalable Brain Atlas geometry for the Allen**

**Mouse Brain Common Coordinate Framework v3 (ABA_v3). Each atlas polygon is filled by the value of the region it represents, resolved by exact acronym match, then by averaging descendant regions present in the data, then by the nearest ancestor present in the data. Participation coefficient is shown on a viridis scale from 0 to 0.8 (a) and WMDz on a diverging blue–white–red scale from −3 to 3 (b), as in Figure 6; two values below −3 are shown at the end of the scale. Polygons are left grey where no value could be resolved. This occurs in three cases: the region lies outside the 198 analyzed grey-matter regions (for example fiber tracts and ventricles); the region was excluded before network construction because its activation count had zero variance or no valid data in that condition and sex; or the region became isolated at the 10% density threshold and was removed from the graph. Grey coverage therefore differs between conditions and between sexes.**

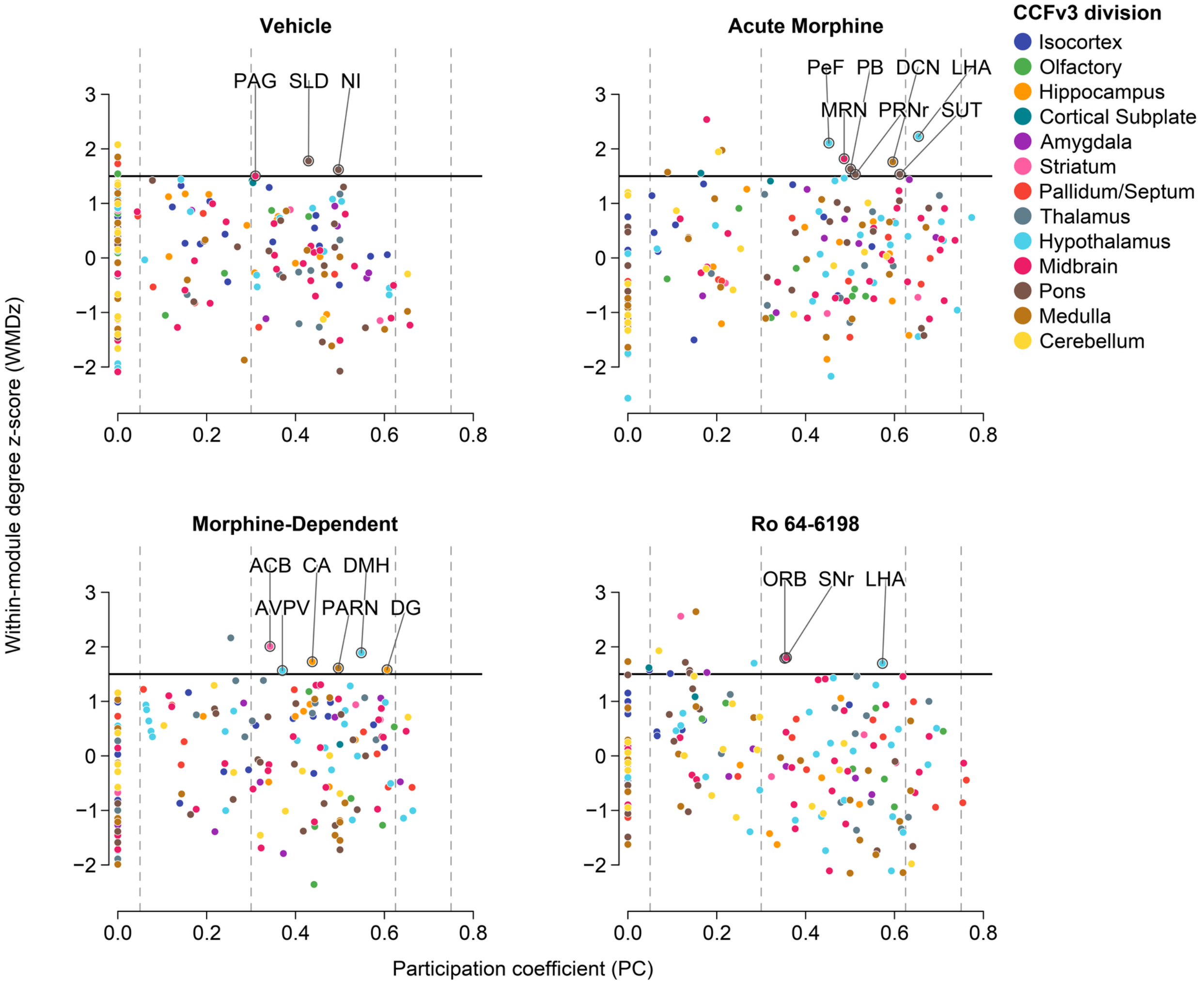


**Figure S3. Cartographic node roles in the combined whole-brain co-activation networks. Guimerà–Amaral cartography for each condition (Vehicle, Acute Morphine, Morphine-Dependent, Ro 64-6198), derived from the combined, sex-pooled co-activation networks (within-sex mean-centered log10 regional activation counts pooled across sexes; n = 12 per condition, n = 10 for Morphine-Dependent; 10% edge-density threshold; consensus-Louvain modules). Each point is one grey-matter region, positioned by its participation coefficient (PC, x-axis) and within-module degree z-score (WMDz, y-axis) and colored by CCFv3 division (CCFv3 division key). The within-module degree z-score expresses how strongly each region is connected to other regions within its own module, in standard deviations relative to the other members of that module; the solid horizontal line marks WMDz = 1.5, the hub threshold (the lower bound of the canonical high-WMDz range; see Supplementary Methods). Regions above this line are hubs within their module. Dashed vertical lines mark the participation-coefficient boundaries of the Guimerà–Amaral role scheme, which subdivides hubs by how widely their connections are distributed across modules. As in Figure 6, only connector hubs (R6; WMDz ≥ 1.5 with intermediate participation, 0.30 < PC ≤ 0.75) are labeled by region abbreviation, whereas provincial hubs (WMDz ≥ 1.5, PC ≤ 0.30) and non-hub nodes are shown but unlabeled. Region abbreviations follow Allen CCFv3 nomenclature.**
